# Discovering and targeting dynamic drugging pockets of the oncogene KRAS-G12D

**DOI:** 10.1101/2022.07.01.498403

**Authors:** Zheyao Hu, Jordi Martí

**Affiliations:** Department of Physics, Polytechnic University of Catalonia-Barcelona Tech, B4-B5 Northern Campus UPC, Barcelona, Catalonia, Spain

## Abstract

Activated KRAS-G12D mutations are the one of most frequent oncogenic drivers in human cancers. Unfortunately, no therapeutic agent directly targeting KRAS-G12D has been clinically approved yet, with such mutated species remaining undrugged. Notably, cofactor Mg^2+^ is closely related to the function of small GTPases, but no investigation has been done yet on Mg^2+^ when associated with KRAS. Herein, through microsecond scale molecular dynamics simulations we have found that Mg^2+^ plays a crucial role in the conformational changes of the KRAS-GDP complex. We have located two brand new druggable dynamic pockets exclusive to KRAS-G12D. Using the structural characteristics of these two dynamic pockets, we designed in silico the inhibitor DBD15-21-22, which can specifically and tightly target KRAS-G12D-GDP-Mg^2+^ ternary complex and we have verified that DBD15-21-22 is harmless for wild-type KRAS. Overall, we provide two brand new druggable pockets located on KRAS-G12D, as well as suitable strategies for KRAS-G12D inhibition.

## 1 Introduction

KRAS proteins play an important role in various cellular processes such as cell proliferation, differentiation, and survival^1, 2^. As a critical hub in cell signalling networks, KRAS function as binary molecular switches^3^: they can be activated by upstream receptors such as EGFR^4–6^ and they can regulate the downstream MAPK^7, 8^, PI3K^9, 10^ and RALGDS–RAL^11, 12^ pathways. KRAS mutations impair both the intrinsic and GAP stimulated GTP hydrolysis activity^13^, leading to hyperactivation of KRAS signalling and ultimately cancer. After near 40 years of efforts, major breakthrough was made for KRAS-G12C inhibition^14^. The covalent binding of designed inhibitors to CYS12 induces a targetable allosteric switch-II pocket, eventually blocking KRAS-G12C^15–18^. Despite these progresses, no therapeutic agent directly targeting the most prevalent and oncogenic KRAS-G12D mutation has been clinically approved yet. Up to now, several strategies have been proposed for targeting KRAS-G12D, such as indole-based inhibitors (BI-2852) of the Switch-I/II pockets^19^, piperazine-based compound TH-Z835 for ASP12^20^, KAL-21404358 for PRO110^21^, a multivalent small molecule 3144 for pan-RAS inhibition^22^ and a three cyclic peptide for GTP-bound KRAS-G12D^23^. However, none of these molecules passed further clinical studies due to lack of precise suitable druggable pockets for inhibition and the picomolar affinity of guanine nucleotide to KRAS^24–26^. Therefore, how to interrupt mutated KRAS-G12D oncogene remains a real challenge both in the clinical and scientific research.

Clinical data have implicated not only that mutated RAS isoforms vary by tissue and cancer types, but also within the most frequently KRAS-mutations the mutated amino acid residues of KRAS are also different for each cancer. For instance, among KRAS mutation predominant cancers, KRAS-G12C mutations mainly occur in lung cancer (45%), whereas KRAS-G12D mutations are related with more than 50% of pancreatic ductal adenocarcinoma cases^27–29^. The KRAS mutations display high tissue-specific abilities to drive tumorigenesis, strongly suggesting that mutation of only one KRAS residue can lead to significant mechanistic changes in KRAS behaviour. This leads to consider that there are usually only molecular scale differences between different KRAS mutants, such as one site GLY12 substitutions. Therefore, the detailed information on atomic interactions and local structures at the all-atom level of KRAS can be of crucial importance for oncogenic KRAS research. Until now, several studies has been published on the effects of G12D mutation on the structure and conformation changes of KRAS^30–34^. However, previous studies mainly focus on oncogenic protein themselves and are less associated with KRAS surroundings, although cancer is closely related to oncoproteins and the surrounding environment^35, 36^. So that more detailed conformations and local structures of KRAS still remain to be revealed.

As one of the important RAS cofactors, the ion Mg^2+^ is coordinated in an octahedral arrangement with a high affinity on RAS proteins^24, 37^. Mg^2+^ has been established essential for both guanine nucleotide binding and GTP-hydrolysis of RAS proteins. For instance, in HRAS the difference in affinity between HRAS and guanine nucleotide in the presence or absence of Mg^2+^ is ∼ 500-fold^38–40^. However, HRAS is rarely mutated in human cancers with only ∼ 10% rate found in bladder and cervical cancers^27–29^. The mechanism of Mg^2+^ interaction with the most prevalent oncogenic KRAS species has not been investigated yet. Herein, through long-time scale molecular dynamics (MD) simulations at all-atom level, we revealed that cofactor Mg^2+^ plays a crucial role in the conformational changes of KRAS. The mutation of GLY12 site in KRAS-G12D triggers a distinct shift in the interaction patterns between Mg^2+^ and KRAS finally generating two druggable exclusive dynamic pockets on the KRAS-G12D-GDP-Mg^2+^ ternary complex surface. With full use of the structural characteristics of these two dynamic pockets we have designed in silico the specific inhibitor DBD15-21-22, a derivative of benzothiadiazine^41^ (DBD), which can specifically and tightly target the KRAS-G12D oncogene stabilising the inactive state of KRAS-G12D-GDP. The reliability of this finding has been validated by subsequent molecular dynamics simulations of KRAS-G12D together with DBD15-21-22 where, by analysing the differences between the three-dimensional structure of DBD15-21-22 and guanine nucleotide and combining them with the molecular dynamics simulation results of DBD15-21-22 and wild-type KRAS, we can propose that DBD15-21-22 will be harmless for wild-type KRAS.

## 2 Results

In this first part of the work, we investigated the conformational changes of the three different isoforms of KRAS in aqueous ionic solution. The only difference between these three KRAS is located at codons 12, with the sequences and initial structures of the three isoforms shown in Figure 1 of “Supporting Information” (SI). Atomic detail sketches of GDP and the main residues described in this part are reported in SI-Figure 2. All meaningful atom-atom distances as a function of time and bond lifetimes have been reported in SI.

**Figure 1:**
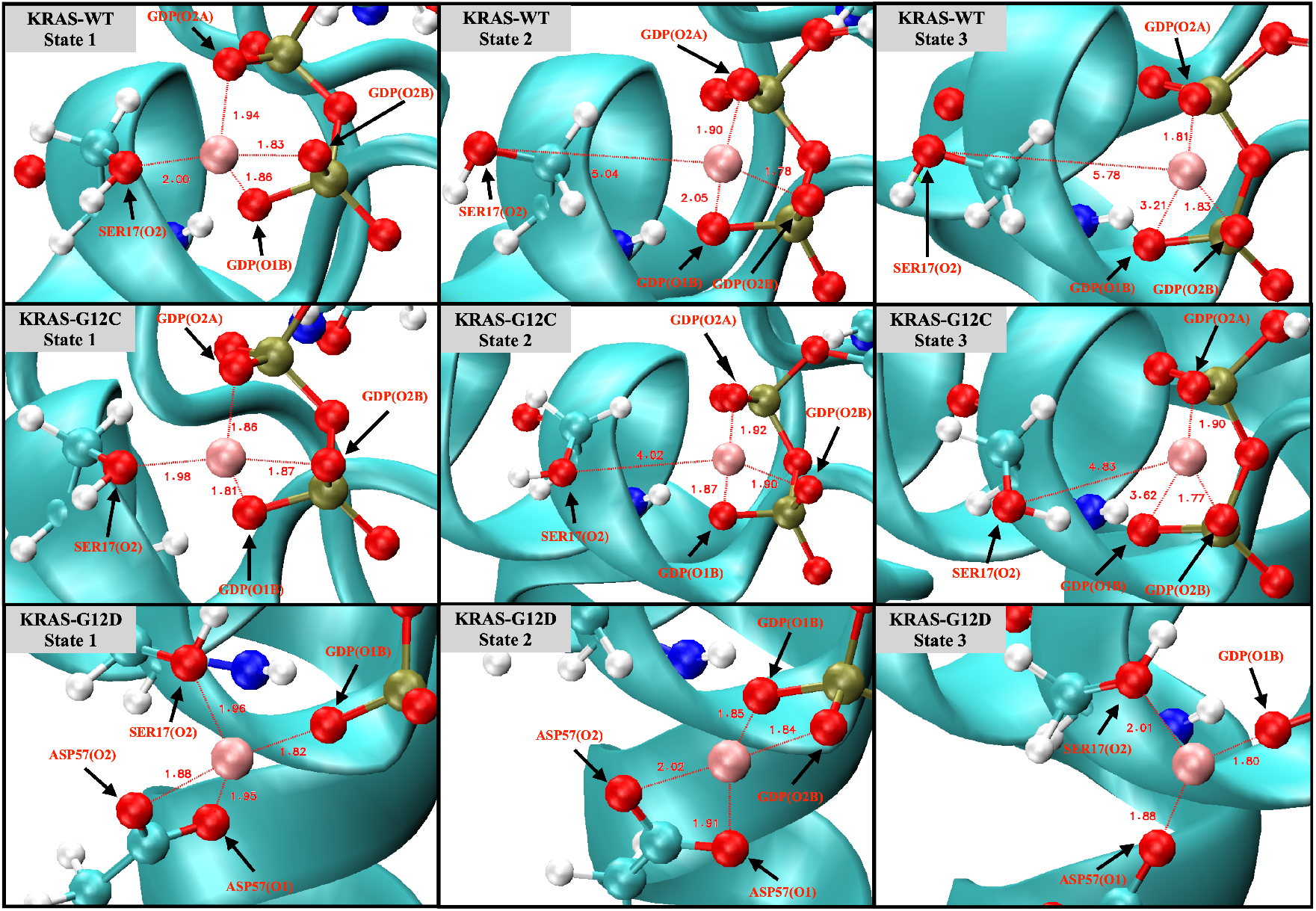
Snapshots of the coordination bonds between Mg^2+^ and SER17, ASP57 and GDP. Water molecules have been hidden for the sake of clarity.

### The fluctuations and stability of KRAS protein conformations with different mutation isoforms

Root Mean Square Deviations (RMSD) and Root Mean Square Fluctuations (RMSF) of KRAS-WT and the two mutant isoforms were firstly analysed (SI-Figure 3A and 3B). Both properties are defined in section (4). RMSD results show the fluctuations and stability of the conformations of the three species. An overall view of the evolution of conformational changes is shown in SI-Figure 3C-3E. We found that for KRAS-WT, there was a distinct conformational fluctuation around 1.9 *µ*s during the simulation. In a similar fashion, for KRAS-G12C a large conformation change was observed after 1 *µ*s. KRAS-G12D exhibited a distinct conformational stability different from that of KRAS-WT and KRAS-G12C, with overall stability and no significant changes. From the perspective of residues, RMSF reveals its flexibility during the full simulation. Switch-I (SW-I) and Switch-II (SW-II) are the regions of the protein which have been recently identified as potential drugging sites^29, 34, 42^ and they show high conformational flexibility compared to other structures of KRAS. Noticeably, we observed that the flexibility of residues in the SW-I domain of KRAS-G12D is around 2-fold smaller than that of KRAS-WT and KRAS-G12C. Combining RMSD and RMSF results, the KRAS-G12D demonstrated significant stability along full 5*µ*s MD simulations. This suggests that (1) the different isoforms of KRAS are not sharing the same mechanisms in conformational changes; and (2) it corroborates that to a large extent, the conformational change of KRAS protein is mainly embodied by SW-I and SW-II. These findings are in overall good agreement with previous computational studies^43^.

### Mutation of GLY12 enhances the interaction between P-loop and SW-II domain

The GLY12 mutation of KRAS protein also shows significant effects on the interaction of P-loop with Switch-II (SI-Figures 4 and 5). GLY12 has weak hydrogen bonding interactions with GLY60 and GLN61, but the *HB* interactions are enhanced when the GLY12 mutated to CYS or ASP. Compared with the GLY12 of KRAS-WT, residue CYS12 contains a sulfhydryl group, so that the HB interaction between CYS12 (H1) and GLN61 (O1) is mildly enhanced (SI-Figure 4B). When GLY12 mutated to ASP, it contains two active HB electron donor sites (carboxyl group). This reverses the surface electrical properties of residue 12, with ASP12 having a longer side chain. The interaction between position 12 and residues GLY60 and GLN61 is significantly enhanced so that they can form 3 simultaneous strong *HB* with GLY60 and GLN61 (SI-Figure 5).

In addition, the mutation of GLY12 also affects the interaction of VAL8 (near the P-loop) with THR58 (located on SW-II) (SI-Figure 6). In KRAS-WT case, the *HB*_*V AL*8(*O*3)−*THR*58(*H*2)_ and *HB*_*V AL*8(*O*3)−*THR*58(*H*3)_ are alternatively generated, but the lifetime of *HB*_*V AL*8(*O*3)−*THR*58(*H*2)_ is slightly longer than that of *HB*_*V AL*8(*O*3)−*THR*58(*H*3)_. In the KRAS-G12C case, the lifetime of *HB*_*V AL*8(*O*3)−*THR*58(*H*2)_ is increased so that such pair remains present for the entire simulation period, while *HB*_*V AL*8(*O*3)−*THR*58(*H*3)_ interactions are mostly eliminated. Furthermore in the case of KRAS-G12D, the lifetimes of both *HB*_*V AL*8(*O*3)−*THR*58(*H*2)_ and *HB*_*V AL*8(*O*3)−*THR*58(*H*3)_ are significantly enhanced, with these two *HB* being present throughout the whole simulation time. Notably, *HB*_*V AL*8(*O*3)−*THR*58(*H*3)_ was more stable than *HB*_*V AL*8(*O*3)−*THR*58(*H*2)_ in KRAS-G12D case, which is a clear difference compared to KRAS-WT and KRAS-G12C.

### Strong coordination interactions between Mg^2+^ and SER17, ASP57, GDP in KRAS-G12D

Noncovalent interactions, such as hydrogen bonds (*HB*) and coordination bonds (*CB*) are important for proteins to maintain their tertiary structure as well as for protein-cofactor interactions. We have analysed the structure of the three different KRAS isoforms in the aqueous ionic solution by time-dependent atomic site–site distances between selected atomic sites, to find out and estimate interactions between Mg^2+^ and KRAS (SI-Figure 7). The full set of averaged values and lifetimes of *HB*/*CB* are also reported in Table 2 of SI. We have selected in Fig. 1 representative snapshots from the full MD trajectory of 5 *µ*s where the pattern and mechanism of formation and breaking of *CB* involving Mg^2+^ is clearly seen. Interestingly, cofactor Mg^2+^ exhibits a unique role in the conformational change of KRAS-G12D, while has little influence on KRAS-WT and KRAS-G12C. As shown in Fig. 1, for the KRAS-G12D case, Mg^2+^ can form *CB* with SER17 (oxygen labelled ‘O2’, see SI Fig.2), ASP57 (O1, O2) of KRAS-G12D and oxygens O1B, O2B of GDP (state 1). During the first 500 ns, 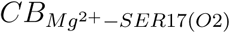 is in a dynamic fluctuation, can be formed and broken (state 2). In the next 4.5 *µ*s, bonds 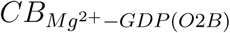 and 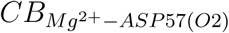 are broken. Finally, Mg^2+^ forms strong *CB* with GDP (O1B), ASP57 (O1) and SER17 (O2) (state 3). In KRAS-WT and KRAS-G12C cases, the interaction patterns of Mg^2+^ with KRAS and GDP are similar each other but they are very different from the KRAS-G12D case. In particular, Mg^2+^ form CB with SER17 (O2) and O2A, O1B, O2B atoms of GDP, while lifetimes of Mg^2+^-SER17 (O2) and Mg^2+^-GDP (O1B) are rather short. Once the 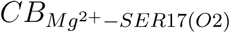 bond was broken, Mg^2+^ was subsequently released from the binding site of KRAS-SER17.

### GDP plays an important regulatory role in the conformational change of SW-I

In addition to the aforementioned *CB*/*HB* interactions in KRAS-GDP-Mg^2+^ complex, the *HB* interactions have been also investigated between SW-I and GDP (SI-Figure 8). We have found that the *HB* interaction of SW-I (ASP30) with GDP is important for the transition of the SW-I conformation. The H2 and H3 atoms of GDP are very sensitive: When they form hydrogen bonds with the oxygen atoms of ASP30, these *HBs* between ASP30 and GDP can limit the conformational change of SW-I within a small range. From the time evolution of selected atom–atom distances *d*(*t*) in SI-Figure 8, we can see that the strength of the *HB* interaction between ASP30 and GDP are KRAS-G12D *>* KRAS-WT *>* KRAS-G12C. This matches the RMSD variation of each KRAS in SI-Figure 3A. Correspondingly, KRAS-G12D exhibits extreme conformational stability during intervals of at least 5 *µ*s of the simulations. We also observed SW-I of KRAS-WT opening largely while the *HB* interaction between ASP30 and GDP breaks around 1.9 *µ*s. Finally, since ASP30 has the weakest *HB* interaction with GDP in the KRAS-G12C case, the latter exhibits large conformational changes around 1 *µ*s.

### The dominant conformations of wild-type and mutated KRAS

Gibbs free energy profiles are of high significance to characterise the dominant conformations of KRAS during simulation. In the present work, such analytical tools will allow us to directly track the effects of GLY12 mutations on the dominant conformations of the KRAS. Since the main movement of the protein is concentrated in SW-I and SW-II and our main research object is KRAS-GDP-Mg^2+^ ternary complex we chose to compute Gibbs free energy landscapes by using two specific variables such as RMSD and radius of gyration (*R*_*g*_), see section 4. The method employed to obtain the free energy profiles has been the so-called “Principal Component Analysis”^44, 45^, where the components RMSD and *R*_*g*_ worked as reaction coordinates (Figure 2). In the KRAS-WT case, we detect two free energy basins (I, II). Basin II is the one with lowest free energy (set to 0 kJ/mol) and it has been chosen as reference so that basin I shows a barrier of 4.6 kJ/mol, when transitions from basin I to II are considered. These two Gibbs free energy basins represent the two main stable states of KRAS-WT during MD simulations. We can obtain from MD trajectories that in state I, SW-I is slightly open while in the dominant basin II SW-I opens largely and SW-II tends to be closer to the P-loop (region containing the mutated codon 12 and depicted in orange colour). Several Gibbs free energy basins were also detected in the KRAS-G12C. In this case, the representative conformation of basin I is similar to the corresponding one in KRAS-WT, while basin II is divided into several small energy basins, with IIa and IIb as examples. The dominant conformation is now located on basin IIb (0 kJ/mol), with basin I at 2.7 kJ/mol and basin IIa at 1.0 kJ/mol. These basins are separated by low free energy barriers and they are moderately easy to access by the system. The differences between basins IIa and IIb are mainly on SW-I, with the SW-I conformation of IIb being more open than that of IIa. The comparison between KRAS-WT, KRAS-G12C and KRAS-G12D shows a clear difference: a single Gibbs free energy basin has been found for KRAS-G12D, what indicates that such species has only one dominant stable conformation. As it can be seen from the representative crystal structure of this particular free energy basin I (Figure 2C), this dominant conformation of KRAS-G12D corresponds to SW-I slightly open, meanwhile the distance between SW-II and P-loop is relatively tight. This is because the strong *CB* interaction of Mg^2+^ with GDP (O1B), SER17 (O2) and ASP57 (O1) can stabilise the distance between SW-II, GDP and the *α*-helix of KRAS, where SER17 is located. We have also observed that the ASP12 shows strong interaction with GLY60 and GLN61, both located at SW-II. At the same time, hydrogen atoms H2 and H3 of GDP can form stable *HB* interactions with ASP30, helping to stabilise the distance between SW-I and GDP.

**Figure 2:**
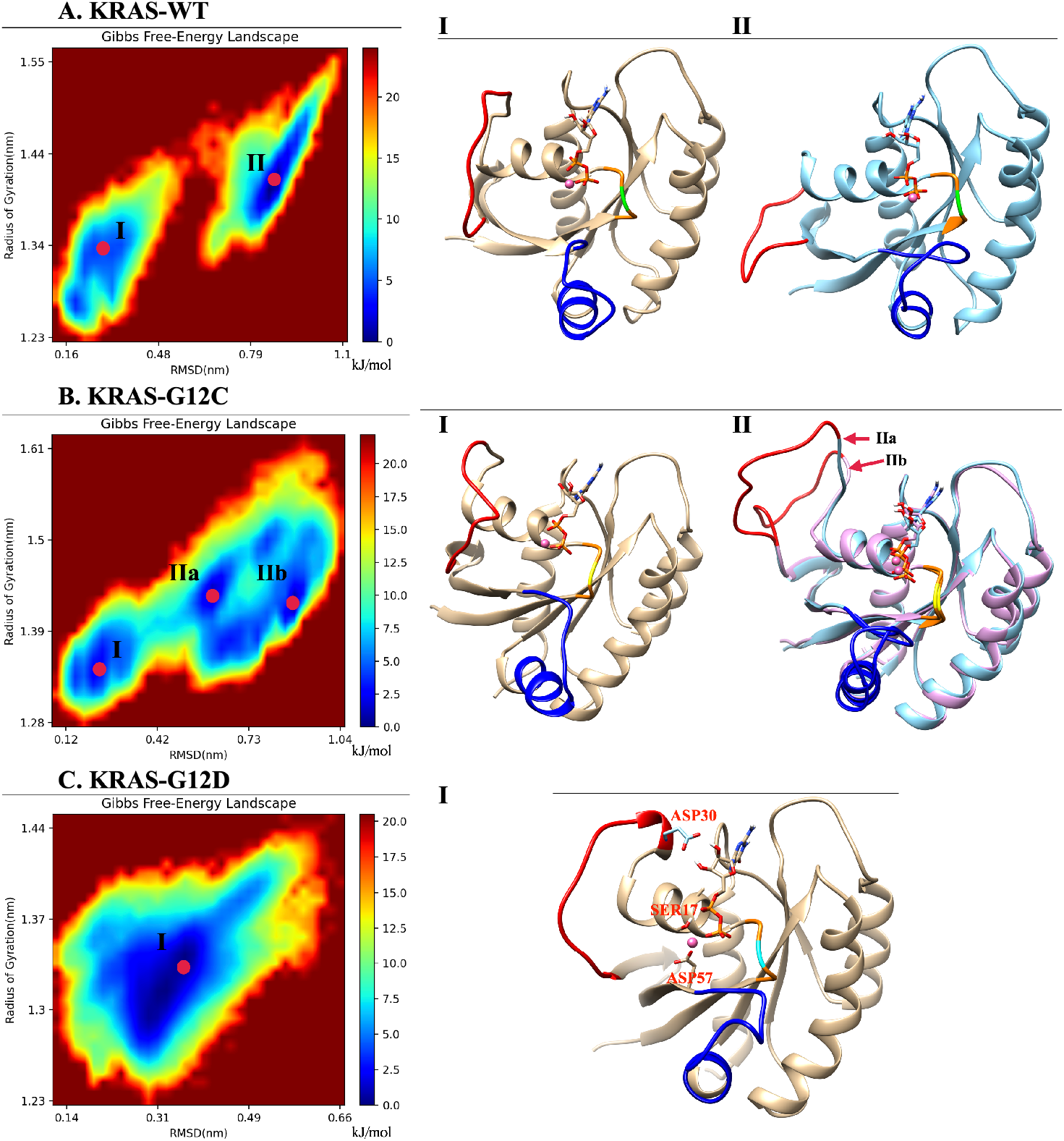
Gibbs free energy landscapes and representative conformations: (A) Landscapes of the KRAS-WT-GDP-Mg^2+^ complex and the representative dominant conformations of basins I and II; (B) Landscapes of KRAS-G12C-GDP-Mg^2+^ complex and the representative dominant conformations of basins I and II; (C) Landscapes of the KRAS-G12D-GDP-Mg^2+^ complex and the single representative dominant conformation (I). Selected groups: backbone atoms of residue 10 to 72; GDP and the Mg^2+^ binding on KRAS. Red: SW-I; Blue: SW-II; Orange: P-loop; Green: GLY12; Yellow: CYS12; Cyan: ASP12.

### The two unique druggable dynamic pockets on KRAS-G12D

From the previous analysis, KRAS-G12D exhibited a stable conformation that was significantly different from those of KRAS-WT and KRAS-G12C and Mg^2+^ exhibited a unique binding mode in KRAS-G12D surface. Mg^2+^ can form strong *CB* interactions with GDP (O1B), SER17 (O2) and ASP57 (O1). Well established experimental results^24, 37^ showed that the octahedral structure is the most stable structure for Mg^2+^ first shell coordination, which is fully consistent with our simulation results. When we take into account the interaction of KRAS-GDP-Mg^2+^ complexes in aqueous Mg-Cl solution, we find the dynamic water pocket I in the Mg^2+^ binding area (left side of Figure 3). We call it “dynamic” in the following sense: In the octahedral structure of Mg^2+^, the fluctuation frequencies of the coordinated water molecules (in positions *H*_2_*O* − 1, *H*_2_*O* − 2 and *H*_2_*O* − 3) are lower than those of free water molecules in solution, i.e., the former remain in their positions for much longer periods of time than the latter (nanoseconds compared to the picosecond time scale for bulk water *HB* dynamics^46^). At the same time, the coordinated water molecules can be exchanged with other water molecules in the solution (SI-Figure 9). Further analysis of the trajectory uncovered another dynamic water pocket II, which is located between the phosphate group of GDP and the ASP12 (right side of Figure 3). This dynamic pocket can accommodate only one *H*_2_*O* molecule, through *HB* interactions of *H*_2_*O* with GDP (O2B) and ASP12 (O1/O2) (SI-Figure 10). Further, the water molecule in this pocket is in a dynamic fast exchange with other water molecules in the solution.

**Figure 3:**
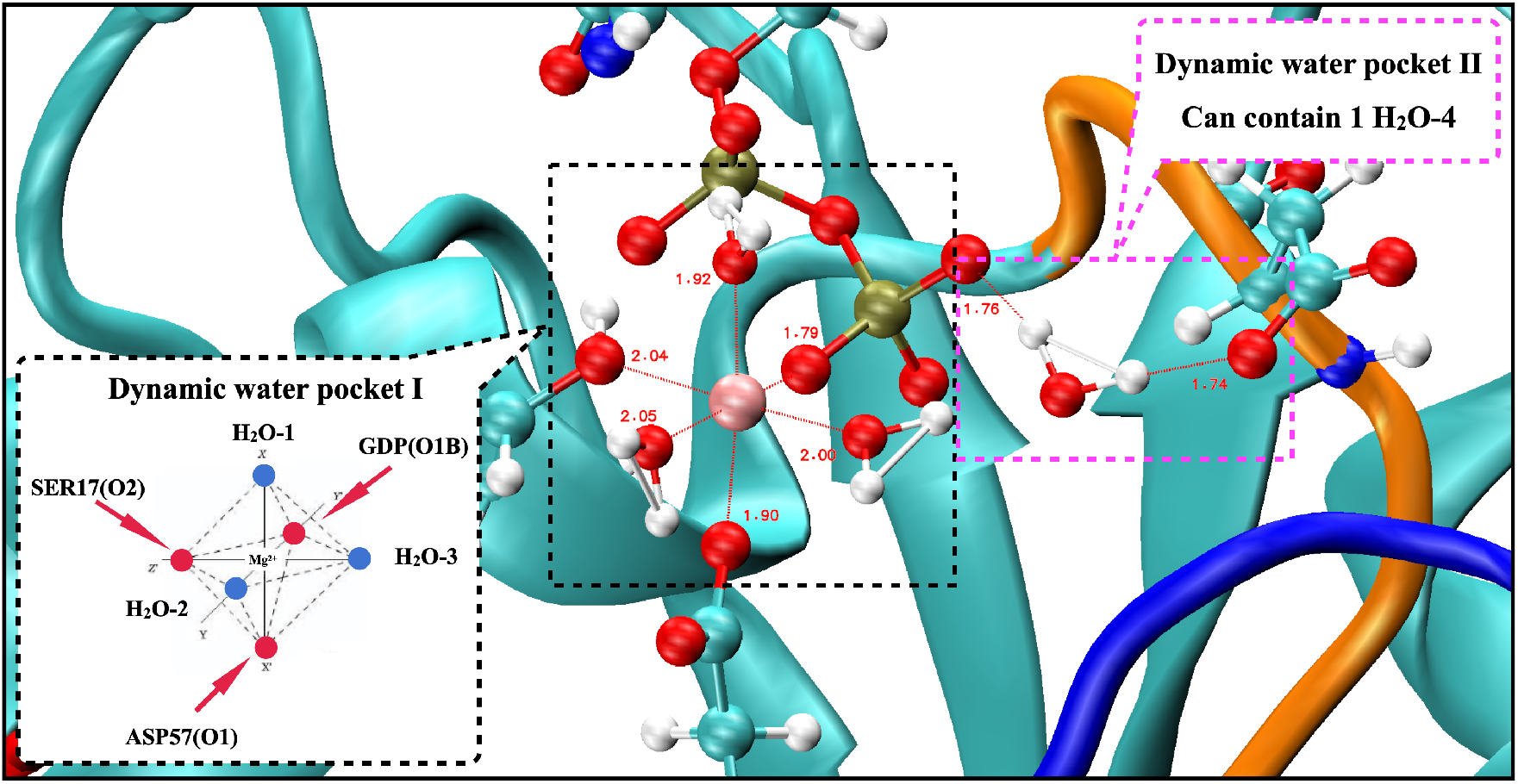
Snapshot of two dynamic pockets on the surface of KRAS-G12D-GDP-Mg^2+^ ternary complex.

### Structure-based drug design and targeting the dynamic water pockets on KRAS-G12D

In the second part of the work, we have designed a drug in silico able to block GDP at its main cavity in a permanent way. There are several identified ways to target KRAS, being the most relevant^47^: inhibition of RAS expression, interference with RAS post-translational modifications, inhibition of RAS function or targeting specific downstream effectors. Furthermore, our strategy is different of the one taken by the designers of recent drug *Sotorasib*^48^. We observed that in recent years, selective binding ligands for Mg^2+^ have gradually become known^49^. In dynamic water pocket I, *H*_2_*O* − 2 and *H*_2_*O* − 3 molecules can exchange with other water molecules in the aqueous solution. The two oxygen atoms of *H*_2_*O* − 2 and *H*_2_*O* − 3 are on the same horizontal plane, and the distance between the two oxygen atoms is comparable to the size of the bidentate ligand. This feature makes possible to design the selective inhibitors that can replace these two water molecules and target this dynamic water pocket I. Herein, we designed a series of inhibitors for these two dynamic water pockets using a benzothiadiazine, specifically 3,4-dihydro-1,2,4-benzothiadiazine-1,1-dioxide (*C*_7_*H*_8_*N*_2_*O*_2_*S*, DBD) as a template. The most suitable species obtained has been designed as DBD15-21-22 (see Figure 4A), since it is a DBD derivative, similar to some of those reported in Ref.^50^.

**Figure 4:**
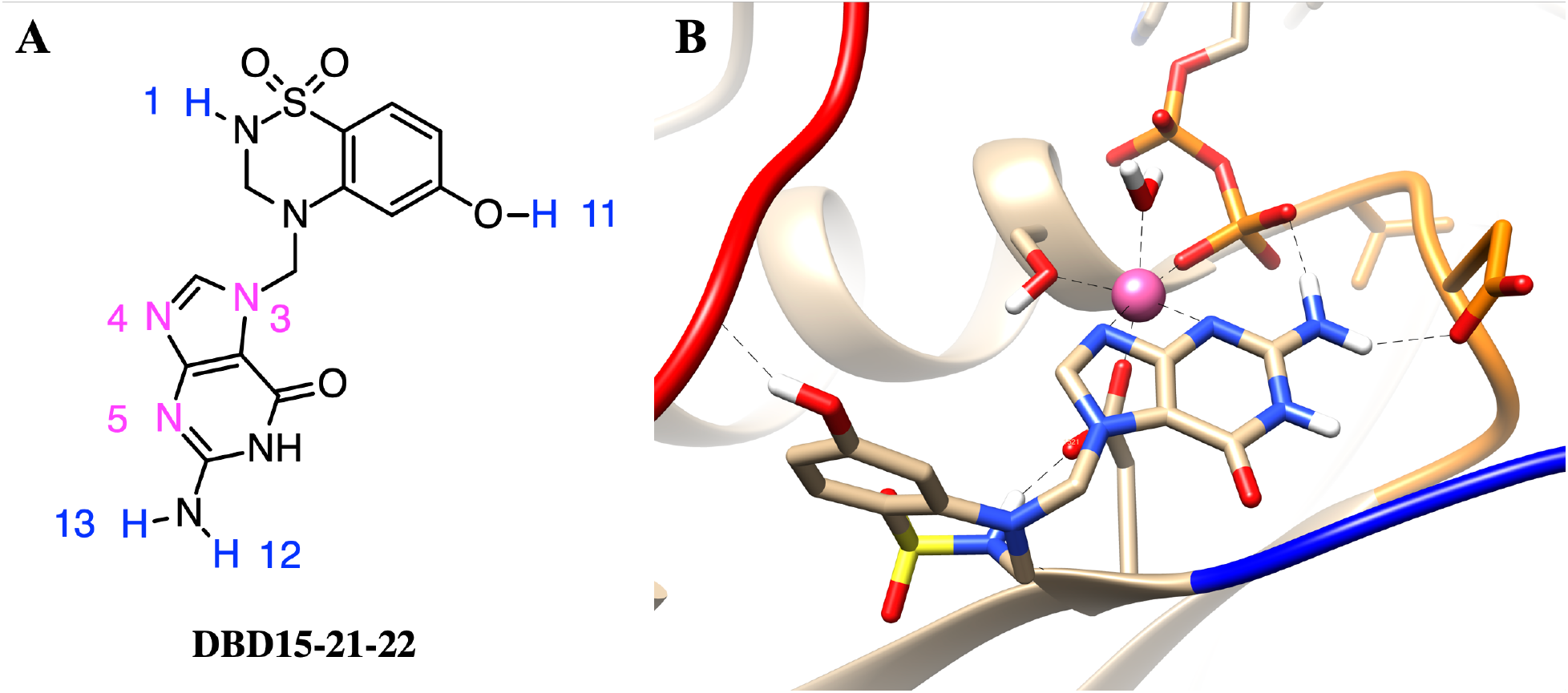
(A) Chemical structure of designed inhibitor DBD15-21-22 and (B) sketch of the interaction of details DBD15-21-22 with KRAS-G12D. Colours as in Figure 2.

### DBD15-21-22 can target KRAS-G12D and bind dynamic water pockets I and II

To further study the structure of DBD15-21-22 interaction with KRAS-G12D, we performed microsecond time scale MD simulations of DBD15-21-22 together with KRAS-G12D. As we can see from the MD results, the DBD15-21-22 is tightly bound to its binding pocket (Figure 4B). The N4 and N5 of DBD15-21-22 can replace *H*_2_*O* − 2 and *H*_2_*O* − 3 in the octahedral structure of Mg^2+^ and then specifically bind to the dynamic water pocket I. Benefiting from the location and size of the amino group (−*NH*_2_) near the N4 and N5, the −*NH*_2_ group can perfectly replace the *H*_2_*O* − 4 in the dynamic water pocket II. These two concerted actions enable DBD15-21-22 to bind tightly and specifically to the two dynamic water pockets on KRAS-G12D-GDP-Mg^2+^ complex. In addition, other active sites in DBD15-21-22 can also form stable *HB* interactions with other residues of KRAS-G12D. In particular (see Figure 4A), H1 atom of DBD15-21-22 can form *HB* with ASP57 (O2) and H11 atom on the phenolic hydroxyl (−*OH*) group can form *HB* with multiple residues on SW-I to lock the conformational change of SW-I. In order to estimate the detailed *HB* and *CB* interactions between DBD15-21-22 and its unique druggable pocket on KRAS-G12D, we display the time evolution of selected atom–atom distances *d*(*t*) in SI-Figures 11 to 15.

### DBD15-21-22 is harmless to KRAS-WT

As we can see in Figure 5A, in the structure of DBD15-21-22 there exist a guanylate moiety but the binding patterns are quite different from GDP/GTP. When GDP/GTP binds to the GDP/GTP binding pocket (Figure 5E), the O3 of guanine goes deep inside the pocket. DBD15-21-22 also has a guanine group, which is similar to GDP/GTP. The difference is that in DBD15-21-22 (Figure 5A), the DBD part is connected to the N3 atom of guanine; in contrast, the *β*-D-ribofuranosyl in the GDP structure is connected to the N4 atom of guanine (Figure 5C). The drug we designed takes advantage of this subtle difference, and prevents the guanine structure in the DBD15-21-22 from binding to the GDP/GTP pocket through the steric hindrance of DBD part in the DBD15-21-22. Moreover, we investigated the three-dimensional structure of DBD15-21-22 and GDP in aqueous solution (Figure 5D and 5F). DBD15-21-22 can form intramolecular hydrogen bonds (Figure 5D), further preventing DBD15-21-22 from binding to the binding site of GDP/GTP on KRAS, and thereby minimising the toxicity to KRAS-WT.

**Figure 5:**
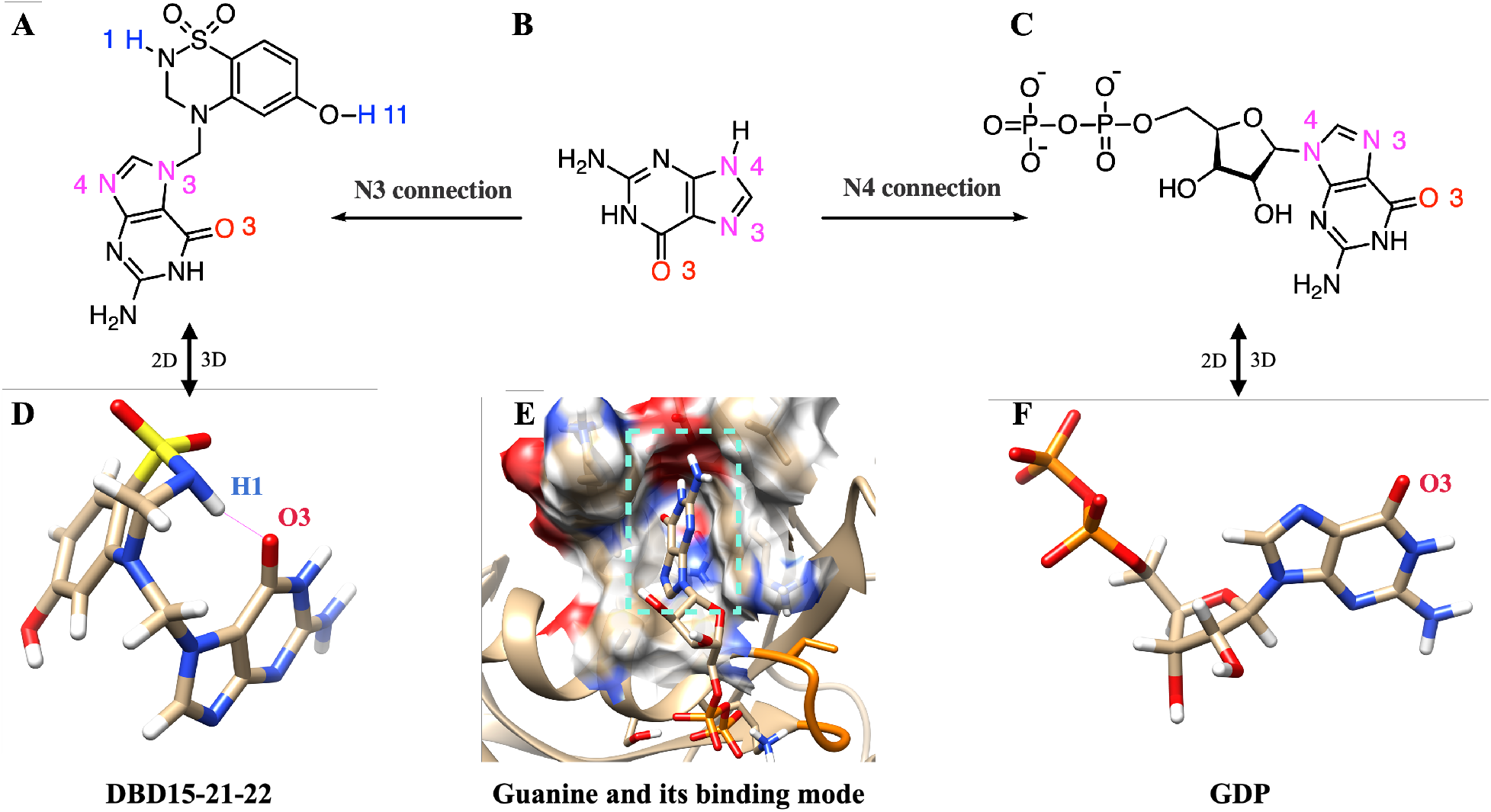
DBD15-21-22 cannot bind to the GDP/GTP binding pocket of KRAS-WT: (A) 2D structure of DBD15-21-22; (B) 2D structure of guanine; (C) 2D structure of GDP; (D) 3D structure of DBD15-21-22 in water; (E) GDP-binding model of KRAS-WT; (F) 3D structure of GDP in water.

To further verify the low toxicity of DBD15-21-22 to KRAS-WT, we also performed MD simulations of DBD15-21-22 together with GDP free KRAS-WT. From the simulation results we can see that DBD15-21-22 was unable to bind to the GDP/GTP binding pocket on KRAS-WT but was dissolved in aqueous solution most of the time (SI-Figure 17A). This finding is further verified by the radial distribution function (RDF) of DBD15-21-22 interaction with aqueous solutions (SI-Figure 17B). We can observed that DBD15-21-22 dissolves well in water, with hydrogen H1 and nitrogen N5 of DBD15-21-22 both forming HBs with *H*_2_*O*. However, in the KRAS-G12D-DBD15-21-22 simulation case, the *HB* interaction of DBD15-21-22 with water was interrupted so that DBD15-21-22 was well bound to the druggable pocket on the surface of KRAS-G12D, as it has been described in Figure 4B).

## 3 Discussion

KRAS mutations are widely present in many cancer cases, but there exist usually only molecular-scale differences between different KRAS mutants, such as in the one site GLY12 mutation. This means the direct information on atomic interactions and local structures at the all-atom level of KRAS can be extremely useful for oncogenic KRAS research. Based on powerful computer simulation tools, we have focused our analysis on the local structure of KRAS proteins when associated with Mg^2+^, GDP, and water. After systematic analysis of meaningful equilibrated data, we can reveal the dynamic differences between KRAS-WT and the mutated species KRAS-G12C and KRAS-G12D at the all-atom level. There exist several important intermolecular interactions affecting the behaviour of KRAS. Most relevant features are: (1) The strong coordination interactions between Mg^2+^, SER17, ASP57 and GDP in KRAS-G12D; (2) The mutation of GLY12 enhances the interaction between P-loop and SW-II domain, with the enhancement magnitude ordered as KRAS-G12D *>* KRAS-G12C; (3) The GLY12 mutation also affects the interaction between GDP and SW-I.

Although GLY12 mutation can enhance the interaction of P-loop with SW-II, in the KRAS-G12C case the interaction of GDP with SW-I was almost totally eliminated, whereas the G12D mutation greatly enhanced the interaction of GDP with SW-I. In Figure 6 we show the alignment of three representative dominant structures of the GDP-bound KRAS-WT, KRAS-G12C and KRAS-G12D. As shown in Figure 6A, the downward sliding of the *α*-helix and *β*-Sheet structures of KRAS-WT promoted the large opening of the SW-I domain. When GLY12 was mutated to ASP12, the interaction between P-loop and SW-II was greatly enhanced, then the relative position of P-loop and SW-II was shortened and stabilised. This promotes strong coordination interactions between Mg^2+^, SER17, ASP57 and GDP, which further stabilises the relative positions of *β*-Sheet and SW-II in the protein structure. Besides, also benefiting from the hydrogen bond interaction between ASP30 and GDP, the fluctuation of SW-I was limited to a small range, which lead to a slightly open stable conformation of SW-I in KRAS-G12D. When GLY12 was mutated to CYS12, although the interaction between P-loop and SW-II was enhanced, it was not enough to stabilise the relative positions of *α*-Helix, *β*-Sheet and SW-II in the structure of KRAS, and the conformational changes were similar to those of the KRAS-WT.

**Figure 6:**
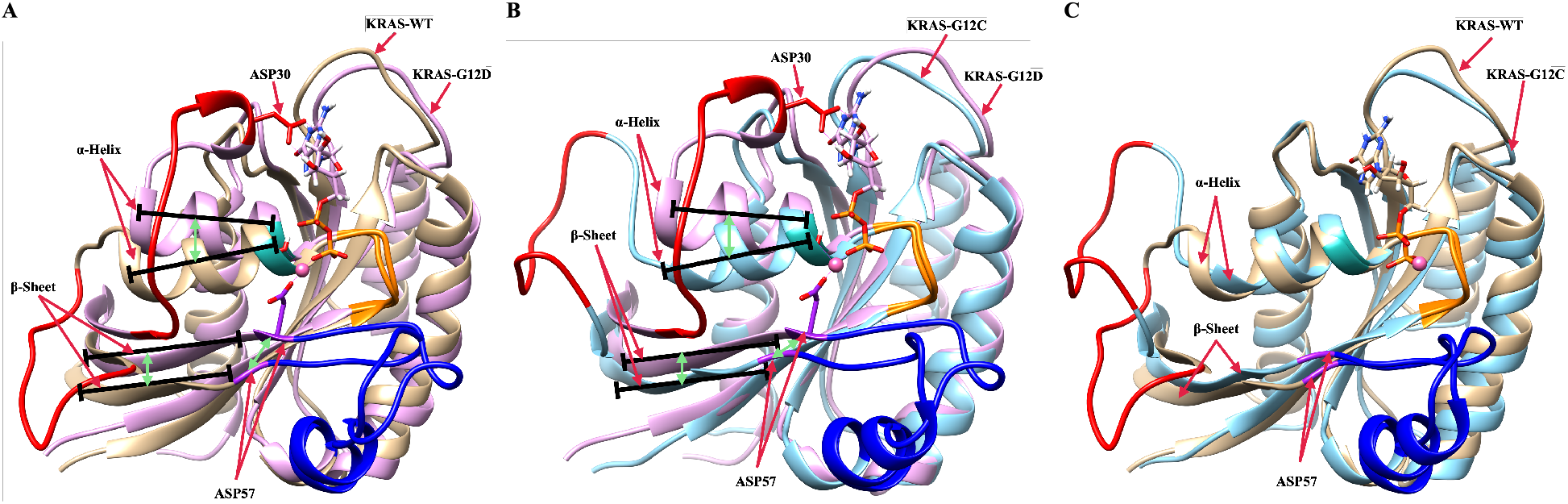
Alignment of three representative dominant structures of the GDP-bound KRAS-WT, KRAS-G12C and KRAS-G12D. (A) Alignment of GDP-bound KRAS-WT and GDP-bound KRAS-G12D; (B) Alignment of GDP-bound KRAS-G12C and GDP-bound KRAS-G12D; (C) Alignment of GDP-bound KRAS-WT and GDP-bound KRAS-G12C.

After deciphering the laws of conformational changes of different KRAS mutants, two drug-gable dynamic water pockets on KRAS-G12D-GDP-Mg^2+^ ternary complex were revealed. We designed a specific inhibitor DBD15-21-22 for these dynamic water pockets, with the aim to lock the KRAS-G12D in the “off” (inactive) state., i.e. when the protein is unable to perform signalling processes and eventually develop cancer. Benefiting from the fact that H1 and O3 atoms of DBD15-21-22 can form intramolecular hydrogen bonds in aqueous solution, we detected no binding effects of DBD15-21-22 towards the GDP/GTP binding pocket of KRAS-WT, whereas our molecular dynamics simulation results show that this KRAS-G12D inhibitor can efficiently bind to the targeting pocket located between SW-I, SW-II and P-loop, thereby efficiently locking KRAS-G12D in the “off” state. Overall, our work provides a new druggable pocket and a reliable protocol for the development of specific inhibitors targeting KRAS-G12D.

## 4 Methods

Our main computational tool has been microsecond scale molecular dynamics. In MD, after the choice of reliable force fields the corresponding Newton’s equations of motion are integrated numerically^51^, allowing us to monitor each individual atom in the system in a wide variety of setups, including liquids at interfaces in solid walls or biological membranes among others ^52, 53^. We fixed the number of particles, the pressure and the temperature of the system, while the volume was adjusted accordingly. MD can model hydrogens at the classical^54^ or quantum levels^55^ and, in addition to energetic and structural properties, MD provides access to time-dependent quantities such as the diffusion coefficients or spectral densities^56^, enhancing its applicability. In the present work we conducted MD simulations of three KRAS isoforms with sequences represented in Fig. 1 of SI. Each system contains one isoform of KRAS-GDP complex fully solvated by 5,697 TIP3P water molecules^57^ in potassium chloride solution at the human body concentration(0.15 M) and magnesium chloride solution concentration(0.03 M) yielding a system size of 19,900 atoms. All MD inputs were generated using CHARMM-GUI solution builder^58–60^ and the CHARMM36m force field^61^ was adopted for KRAS-GDP-Mg^2+^ interactions. The force field used also includes the parameterisation of the species GDP (it can be searched as “GDP” in the corresponding CHARMM36m topology file: https://www.charmm-gui.org/?doc=archive & lib=csml) All bonds involving hydrogens were set to fixed lengths, allowing fluctuations of bond distances and angles for the remaining atoms. Crystal structure of GDP-bound KRAS proteins was downloaded from RCSB PDB Protein Data Bank^62^, file name “4obe”. The three sets of KRAS-GDP complex(wild type, G12C mutate and G12D mutate) were solvated in a water box, all systems were energy minimised for 50,000 steps and well equilibrated (NVT equilibration in Figure 17 of SI) for 250 ps before generating the production MD. Production runs were performed with an NPT ensemble for 5 *µ*s. The pressure and temperature were set at 1 atm and 310.15 K respectively, in order to simulate the human environment. In all MD simulations, the GROMACS/2021 package was employed^63^. Time steps of 2 fs were used in all production simulations and the particle mesh Ewald method with Coulomb radius of 1.2 nm was employed to compute long ranged electrostatic interactions. The cutoff for Lennard-Jones interactions was set to 1.2 nm. Pressure was controlled by a Parrinello-Rahman piston with damping coefficient of 5 ps^−1^ whereas temperature was controlled by a Nose’
s-Hoover thermostat with a damping coefficient of 1 ps^−1^. Periodic boundary conditions in three directions of space have been taken. We employed the “gmx-sham” tool of the GROMACS/2021 package to performed the Gibbs free energy landscape analysis. The designed inhibitor DBD15-21-22 was parameterised using CgenFF^64, 65^ to get the CHARMM-compatible topology and parameter files. Further, we switched to run another four 1 *µ*s of MD simulations to study the interaction of the inhibitor DBD15-21-22 with KRAS-G12D and to verify the DBD-15-21-22 do harmless on the wild-type KRAS. These four new MD simulations have the same setup as the previous ones. Moreover, the software VMD^66^ and UCSF Chimera^67^ were used for trajectory analysis and visualisation.

The radius of gyration (*R*_*g*_), used as a reaction coordinate in the computation of Gibbs free energy landscapes is defined in Eq.1 as

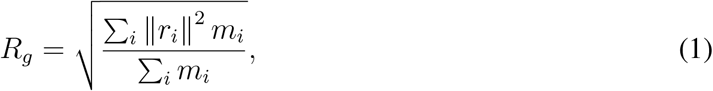

where *m*_*i*_ is the mass of atom *i* and *r*_*i*_ the position of the same atom with respect to the centre of mass of the selected group. Root Mean Square Deviations (RMSD) are defined by Eq.(2)

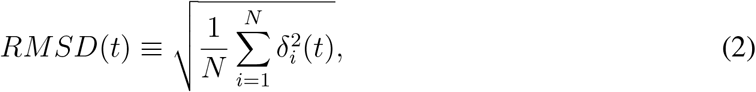

where *δ*_*i*_ is the difference in distance between the atom *i* (located at *x*_*i*_(*t*)) of the catalytic domain and the equivalent location in the crystal structure and Root Mean Square Fluctuations (RMSF) are defined by Eq.(3):

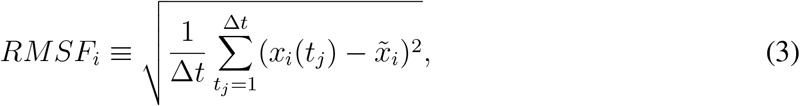

where 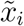 is the time average of *x*_*i*_ and Δ*t* is the time interval where the average has been taken.

## References

1. Cherfils, J. & Zeghouf, M. Regulation of small gtpases by gefs, gaps, and gdis. Physiological reviews 93, 269–309 (2013).

2. Ostrem, J. M. & Shokat, K. M. Direct small-molecule inhibitors of kras: from structural insights to mechanism-based design. Nature reviews Drug discovery 15, 771–785 (2016).

3. Vetter, I. R. & Wittinghofer, A. The guanine nucleotide-binding switch in three dimensions. Science 294, 1299–1304 (2001).

4. Normanno, N. et al. Implications for kras status and egfr-targeted therapies in metastatic crc. Nature reviews Clinical oncology 6, 519–527 (2009).

5. Ardito, C. M. et al. Egf receptor is required for kras-induced pancreatic tumorigenesis. Cancer cell 22, 304–317 (2012).

6. Navas, C. et al. Egf receptor signaling is essential for k-ras oncogene-driven pancreatic ductal adenocarcinoma. Cancer cell 22, 318–330 (2012).

7. Wood, K. W., Sarnecki, C., Roberts, T. M. & Blenis, J. ras mediates nerve growth factor receptor modulation of three signal-transducing protein kinases: Map kinase, raf-1, and rsk. Cell 68, 1041–1050 (1992).

8. Howe, L. R. et al. Activation of the map kinase pathway by the protein kinase raf. Cell 71, 335–342 (1992).

9. Sjölander, A., Yamamoto, K., Huber, B. E. & Lapetina, E. G. Association of p21ras with phosphatidylinositol 3-kinase. Proceedings of the National Academy of Sciences 88, 7908–7912 (1991).

10. Rodriguez-Viciana, P. et al. Phosphatidylinositol-3-oh kinase direct target of ras. Nature 370, 527–532 (1994).

11. Hofer, F., Fields, S., Schneider, C. & Martin, G. S. Activated ras interacts with the ral guanine nucleotide dissociation stimulator. Proceedings of the National Academy of Sciences 91, 11089–11093 (1994).

12. Spaargaren, M. & Bischoff, J. R. Identification of the guanine nucleotide dissociation stimulator for ral as a putative effector molecule of r-ras, h-ras, k-ras, and rap. Proceedings of the National Academy of Sciences 91, 12609–12613 (1994).

13. Hall, B. E., Bar-Sagi, D. & Nassar, N. The structural basis for the transition from ras-gtp to ras-gdp. Proceedings of the National Academy of Sciences 99, 12138–12142 (2002).

14. Ostrem, J. M., Peters, U., Sos, M. L., Wells, J. A. & Shokat, K. M. K-ras (g12c) inhibitors allosterically control gtp affinity and effector interactions. Nature 503, 548–551 (2013).

15. Patricelli, M. P. et al. Selective inhibition of oncogenic kras output with small molecules targeting the inactive state. Cancer discovery 6, 316–329 (2016).

16. Lito, P., Solomon, M., Li, L.-S., Hansen, R. & Rosen, N. Allele-specific inhibitors inactivate mutant kras g12c by a trapping mechanism. Science 351, 604–608 (2016).

17. Janes, M. R. et al. Targeting kras mutant cancers with a covalent g12c-specific inhibitor. Cell 172, 578–589 (2018).

18. Canon, J. et al. The clinical kras (g12c) inhibitor amg 510 drives anti-tumour immunity. Nature 575, 217–223 (2019).

19. Kessler, D. et al. Drugging an undruggable pocket on kras. Proceedings of the National Academy of Sciences 116, 15823–15829 (2019).

20. Mao, Z. et al. Kras (g12d) can be targeted by potent inhibitors via formation of salt bridge. Cell Discovery 8, 1–14 (2022).

21. Feng, H. et al. K-rasg12d has a potential allosteric small molecule binding site. Biochemistry 58, 2542–2554 (2019).

22. Welsch, M. E. et al. Multivalent small-molecule pan-ras inhibitors. Cell 168, 878–889 (2017).

23. Zhang, Z. et al. Gtp-state-selective cyclic peptide ligands of k-ras (g12d) block its interaction with raf. ACS central science 6, 1753–1761 (2020).

24. John, J. et al. Kinetic and structural analysis of the mg (2+)-binding site of the guanine nucleotide-binding protein p21h-ras. Journal of Biological Chemistry 268, 923–929 (1993).

25. Ford, B. et al. Characterization of a ras mutant with identical gdp-and gtp-bound structures. Biochemistry 48, 11449–11457 (2009).

26. Traut, T. W. Physiological concentrations of purines and pyrimidines. Molecular and cellular biochemistry 140, 1–22 (1994).

27. Forbes, S. A. et al. Cosmic: mining complete cancer genomes in the catalogue of somatic mutations in cancer. Nucleic acids research 39, D945–D950 (2010).

28. Prior, I. A., Lewis, P. D. & Mattos, C. A comprehensive survey of ras mutations in cancer. Cancer research 72, 2457–2467 (2012).

29. Moore, A. R., Rosenberg, S. C., McCormick, F. & Malek, S. Ras-targeted therapies. Nat. Rev. Drug Discov. (2021).

30. Lu, S., Jang, H., Nussinov, R. & Zhang, J. The structural basis of oncogenic mutations g12, g13 and q61 in small gtpase k-ras4b. Scientific reports 6, 1–15 (2016).

31. Prakash, P., Hancock, J. F. & Gorfe, A. A. Binding hotspots on k-ras: Consensus ligand binding sites and other reactive regions from probe-based molecular dynamics analysis. Proteins: Structure, Function, and Bioinformatics 83, 898–909 (2015).

32. Chen, C.-C. et al. Computational analysis of kras mutations: implications for different effects on the kras p. g12d and p. g13d mutations. PloS one 8, e55793 (2013).

33. Lu, H. & Martí, J. Long-lasting salt bridges provide the anchoring mechanism of oncogenic kirsten rat sarcoma proteins at cell membranes. The Journal of Physical Chemistry Letters 11, 9938–9945 (2020).

34. Lu, H. & Martí, J. Predicting the conformational variability of oncogenic gtp-bound g12d mutated kras-4b proteins at zwitterionic model cell membranes. Nanoscale 14, 3148–3158 (2022).

35. Whiteside, T. The tumor microenvironment and its role in promoting tumor growth. Oncogene 27, 5904–5912 (2008).

36. Kessenbrock, K., Plaks, V. & Werb, Z. Matrix metalloproteinases: regulators of the tumor microenvironment. Cell 141, 52–67 (2010).

37. Bock, C. W., Kaufman, A. & Glusker, J. P. Coordination of water to magnesium cations. Inorganic Chemistry 33, 419–427 (1994).

38. Tucker, J. et al. Expression of p21 proteins in escherichia coli and stereochemistry of the nucleotide-binding site. The EMBO Journal 5, 1351–1358 (1986).

39. Hall, A. & Self, A. J. The effect of mg2+ on the guanine nucleotide exchange rate of p21n-ras. Journal of Biological Chemistry 261, 10963–10965 (1986).

40. Feuerstein, J., Goody, R. S. & Wittinghofer, A. Preparation and characterization of nucleotide-free and metal ion-free p21 “apoprotein”. Journal of Biological Chemistry 262, 8455–8458 (1987).

41. Novello, F. C. & Sprague, J. M. Benzothiadiazine dioxides as novel diuretics. Journal of the American Chemical Society 79, 2028–2029 (1957).

42. Nussinov, R., Tsai, C.-J. & Jang, H. Oncogenic ras isoforms signaling specificity at the mem-brane. Cancer research 78, 593–602 (2018).

43. Vatansever, S., Erman, B. & Gümüş, Z. H. Oncogenic g12d mutation alters local conformations and dynamics of k-ras. Scientific reports 9, 1–13 (2019).

44. Stein, S. A. M., Loccisano, A. E., Firestine, S. M. & Evanseck, J. D. Principal components analysis: a review of its application on molecular dynamics data. Annual Reports in Computational Chemistry 2, 233–261 (2006).

45. Jolliffe, I. T. & Cadima, J. Principal component analysis: a review and recent developments. Philosophical Transactions of the Royal Society A: Mathematical, Physical and Engineering Sciences 374, 20150202 (2016).

46. Marti, J. Dynamic properties of hydrogen-bonded networks in supercritical water. Physical Review E 61, 449 (2000).

47. Tomasini, P., Walia, P., Labbe, C., Jao, K. & Leighl, N. B. Targeting the kras pathway in non-small cell lung cancer. The oncologist 21, 1450–1460 (2016).

48. Lietman, C. D., Johnson, M. L., McCormick, F. & Lindsay, C. R. More to the ras story: Krasg12c inhibition, resistance mechanisms, and moving beyond krasg12c. American Society of Clinical Oncology Educational Book 42, 1–13 (2022).

49. Walter, E. R., Hogg, C., Parker, D. & Williams, J. G. Designing magnesium-selective ligands using coordination chemistry principles. Coordination Chemistry Reviews 428, 213622 (2021).

50. Hu, Z. & Marti, J. In silico drug design of benzothiadiazine derivatives interacting with phospholipid cell membranes. Membranes 12, 331 (2022).

51. Frenkel, D. & Smit, B. Understanding molecular simulation: from algorithms to applications, vol. 1 (Elsevier, 2001).

52. Nagy, G., Gordillo, M., Guàrdia, E. & Martí, J. Liquid water confined in carbon nanochannels at high temperatures. The Journal of Physical Chemistry B 111, 12524–12530 (2007).

53. Marrink, S. J. et al. Computational modeling of realistic cell membranes. Chemical reviews 119, 6184–6226 (2019).

54. Padro, J., Marti, J. & Guardia, E. Molecular dynamics simulation of liquid water at 523 k. Journal of Physics: Condensed Matter 6, 2283 (1994).

55. Calero, C., Martí, J. & Guàrdia, E. 1h nuclear spin relaxation of liquid water from molecular dynamics simulations. The Journal of Physical Chemistry B 119, 1966–1973 (2015).

56. Martí, J., Padró, J. & Guardia, E. Computer simulation of molecular motions in liquids: Infrared spectra of water and heavy water. Molecular Simulation 11, 321–336 (1993).

57. Jorgensen, W. L., Chandrasekhar, J., Madura, J. D., Impey, R. W. & Klein, M. L. Comparison of simple potential functions for simulating liquid water. J. Chem. Phys. 79, 926–935 (1983).

58. Jo, S., Kim, T., Iyer, V. G. & Im, W. Charmm-gui: a web-based graphical user interface for charmm. Journal of Computational Chemistry 29, 1859–1865 (2008).

59. Brooks, B. R. et al. Charmm: the biomolecular simulation program. Journal of computational chemistry 30, 1545–1614 (2009).

60. Lee, J. et al. Charmm-gui input generator for namd, gromacs, amber, openmm, and charmm/openmm simulations using the charmm36 additive force field. Journal of chemical theory and computation 12, 405–413 (2016).

61. Huang, J. & MacKerell Jr, A. D. Charmm36 all-atom additive protein force field: Validation based on comparison to nmr data. Journal of computational chemistry 34, 2135–2145 (2013).

62. Kouranov, A. et al. The rcsb pdb information portal for structural genomics. Nucleic acids research 34, D302–D305 (2006).

63. Berendsen, H. J., van der Spoel, D. & van Drunen, R. Gromacs: a message-passing parallel molecular dynamics implementation. Computer Physics Communications 91, 43–56 (1995).

64. Vanommeslaeghe, K. & MacKerell Jr, A. D. Automation of the charmm general force field (cgenff) i: bond perception and atom typing. Journal of chemical information and modeling 52, 3144–3154 (2012).

65. Vanommeslaeghe, K., Raman, E. P. & MacKerell Jr, A. D. Automation of the charmm general force field (cgenff) ii: assignment of bonded parameters and partial atomic charges. Journal of chemical information and modeling 52, 3155–3168 (2012).

66. Humphrey, W., Dalke, A. & Schulten, K. Vmd: visual molecular dynamics. Journal of molecular graphics 14, 33–38 (1996).

67. Pettersen, E. F. et al. Ucsf chimera—a visualization system for exploratory research and analysis. Journal of computational chemistry 25, 1605–1612 (2004).

