## Supplemental Information for "Discovering and targeting dynamic drugging pockets of the oncogene KRAS-G12D"

### Supplementary information for: Discovering and targeting dynamic drugging pockets of the oncogene KRAS-G12D

#### ABSTRACT

[1] Department of Physics, Polytechnic University of Catalonia-Barcelona Tech, B4-B5 Northern Campus UPC, Barcelona, Catalonia, Spain

#### 1 Supplementary Notes

##### 1.1 Initial setups and data for molecular dynamics simulations

The three classes of KRAS proteins that have been considered in this work are reported in Fig. 1, where the initial structure of KRAS-WT, was acquired from PDB bank (4obe.pdb). We also show the octahedral structure of $Mg^{2+}$ in the crystal structure, consistent with relevant experimental studies[1,2]. In order to make clear the atomic sites that will be described and analysed in the article, we represent sketches of GDP and the main amino acid structures analysed in the present work in Fig. 2.

In Fig. 3, and from the RMSD values as a function of time, we can see that the conformation of KRAS-G12D is stable during the whole simulation process (5 $\mu$s). The difference from KRAS-G12D is that the conformation of KRAS-WT protein changes greatly in about 1.9 $\mu$s, and the conformation of KRAS-G12C protein changes greatly in about 1 $\mu$s. The RMSF results show the flexibility of amino acid residues in the KRAS structure. From the RMSF results, it can be seen that the structurally active regions of KRAS protein are mainly Switch-I (SW-I) and Switch-II (SW-II) domains. Notably, the flexibility of amino acid residues in the SW-I domain of KRAS-G12D is around 2-fold smaller than that of KRAS-WT and KRAS-G12C.

##### 1.2 Atomic distances and KRAS conformations

Throughout the work it is very important to make sure that binding connections between specific atomic sites are stable enough to be considered as meaningful interactions so that monitoring the instantaneous distances between tagged species is of crucial importance. In our case, the computation of radial distribution functions is not suitable in most cases, given the specificity of the atomic sites involved in the relevant bindings between ions, amino acids and drugs. The series of distances has been reported in Figures 4-16, together with snapshots corresponding to the configurations of highest biophysical relevance. We mainly monitor hydrogens bonds (HB) and coordination bonds (CB).

For instance, in Fig. 6 we compare the effect of different mutations on the interactions between amino acid residues (VAL8-THR58) in the KRAS protein. In KRAS-WT, the HB interaction can last $\sim$2 $\mu$s. When the GLY12 mutated to CYS, the HB interaction between VAL8 and THR58 have been enhanced, interestingly, VAL8 (O3) prefers to form hydrogen bonds with THR58 (H2). In KRAS-G12D case, the hydrogen bond interaction between VAL8 and THR58 is significantly enhanced compared to KRAS-WT and KRAS-G12C. In Fig. 7 the coordination bond analysis of $Mg^{2+}$ is shown. In KRAS-WT case, $Mg^{2+}$ cannot form a CB with ASP57; in the first $\sim$550 ns, $Mg^{2+}$ can form CB with SER17 (O2) and GDP (O2A,O1B,O2B) and the $CB_{Mg^{2+}-SER17(O2)}$ and $CB_{Mg^{2+}-GDP(O1B)}$ are broken, while the $CB_{Mg^{2+}-GDP(O2A)}$ and $CB_{Mg^{2+}-GDP(O2B)}$ are very stable. In KRAS-G12C case, the situation is similar to KRAS-WT. The only difference is that the covalent interaction between $Mg^{2+}$ and SER17 (O2)/GDP (O1B) can last $\sim$700 ns. Finally, in the KRAS-G12D case, the difference are that $Mg^{2+}$ can form stable CB interaction with ASP57, SER17 and GDP (O1B). In the first $\sim$500 ns, the coordination interactions between $Mg^{2+}$, SER17, ASP57 and GDP fluctuated, and then $Mg^{2+}$ formed stable coordination bonds with SER17 (O2), ASP57 (O1) and GDP (O1B). From the selected atom-atom distance data of Fig. 8 we can see that the structure of KRAS-G12C is similar to that of KRAS-WT, with the hydrogen bond interaction between GDP and ASP30 being very weak. However, when GLY12 is mutated to ASP12, the interaction between GDP and ASP30 has been significantly enhanced.

The dynamic change law of three water molecules (in positions $H_2O-1$, $H_2O-2$ and $H_2O-3$) in the dynamic water pocket I is similar, so in Fig. 9 we take the dynamic change law of water molecules at position $H_2O-1$ as an example. From the selected atom-atom distance data, we can see that the exchange period between water molecules in dynamic water pocket I and solution water molecules is $\sim$2.5 $\mu$s. The dynamic water pocket II is consists of the oxygen atom (O2B) of GDP and the oxygen atoms of ASP12 (O1,O2), as shown in Fig. 10 where in left side we show the representative snapshot of dynamic water pocket II and in the right side the distance between ASP12 (O1,O2) and GDP (O2B). As shown in the left side of Fig. 10, the dynamic water pocket II can accommodate one water molecule. And this water molecular has a high exchange frequency with the water molecule in the aqueous solution, with an exchange period of about $\sim$200 ps. Therefore, the time evolution of atom-atom distance $d(t)$ corresponding to the water molecule and GDP/ASP12 is not calculated.

Interestingly, some HB can switch: In Fig. 11 the hydrogen atoms (H12) of DBD15-21-22 can form stable hydrogen bonds interaction with the oxygen atoms (O1,O2) of ASP12. And the ASP12(O1, O2) alternately form stable hydrogen bonds with DBD15-21-22(H12) (see Fig. 11B). Conversely, in Fig. 12B, the hydrogen atoms (H13) of DBD15-21-22 can form stable hydrogen bonds interaction with the oxygen atoms (O2B) of GDP.

The dynamics of the designed drug DBD15-21-22 (H1) indicates (Fig. 13B) that at the beginning of the trajectory it is very far from the ASP57(O2) but after 50 ns it becomes close to the ASP57 (O2) and capable to form stable hydrogen bonds interactions. Further, as shown in Fig. 14B, DBD15-21-22 (H11) can form hydrogen bonds with the oxygen atoms (THR35 (O1), ILE36 (O1) and GLU37 (O3)) of three different SW-I residues. Finally, from the distance between H1 and O3 atoms in DBD15-21-22, we can find that DBD15-21-22 can form intramolecular hydrogen bonds $HB_{H1-O3}$ when DBD15-21-22 is in aqueous solution.

##### 1.3 Convergence of the equilibrated simulations

Our simulation systems' equilibration input generation option is "NVT Ensemble" and the temperature we set is 310.15K. According to the temperature data we obtained for the equilibration step, our systems were well equilibrated before the production step. as it can be observed from Fig. 18 (three species of KRAS in aqueous solution), Fig. 19 (KRAS-WT and KRAS-G12D with the drug DBD15-21-22 in aqueous solution) and Fig. 20 (the drug DBD-15-21-22 in aqueous solution, and GDP in aqueous solution). Analog figures for total energies in the same system setups have not been included.

#### 2 Supplementary Tables

**Table 1.** Amino acid components of the KRAS proteins. Abbreviations as used in the text and in several figures.

| Full name | Abbreviation |
|---|---|
| Aspartate | ASP |
| Cysteine | CYS |
| Glutamine | GLN |
| Glycine | GLY |
| Lysine | LYS |
| Serine | SER |
| Threonine | THR |
| Valine | VAL |
| Proline | PRO |

**Table 2.** Estimated lifetimes (in ns) of HB and CB interactions (Section 2 of main text, corresponding to Figures 4-8). Total simulation time of 5000 ns. We indicate in "Bond" the two amino acids or ion involved in the HB or CB and the corresponding atoms (hydrogen, oxygen) in parentheses. Double labels indicate averaged values.

| Bond (ion/amino acid (atom)) | KRAS-WT | KRAS-G12C | KRAS-G12D |
|---|---|---|---|
| GLY12 (H3)-GLY60 (O3) | 1 | - | - |
| GLY12 (H3)-GLN61 (O1) | 14 | - | - |
| CYS12 (H1)-GLY60 (O3) | - | 1 | - |
| CYS12 (H1)-GLN61 (O1) | - | 16 | - |
| ASP12 (O1,O2)-GLY60 (H3) | - | - | 683 |
| ASP12 (O1,O2)-GLN61 (H1,H2) | - | - | 51 |
| ASP12 (O1,O2)-GLN61 (H3) | - | - | 225 |
| VAL8 (O3)-THR58 (H2) | 437 | 2100 | 843 |
| VAL8 (O3)-THR58 (H3) | 371 | 10 | 2805 |
| Mg-ASP57 (O1) | - | - | 4991 |
| Mg-ASP57 (O2) | - | - | 497 |
| Mg-SER17 (O2) | 503 | 657 | 4258 |
| Mg-GDP (O2A) | 5000 | 5000 | - |
| Mg-GDP (O1B) | 568 | 689 | 5000 |
| Mg-GDP (O2B) | 5000 | 5000 | 420 |
| ASP30 (O1,O2)-GDP (H2,H3) | 130 | 3 | 1423 |
| ASP30 (O3)-GDP (H2,H3) | 60 | 4 | 95 |

**Table 3.** Estimated lifetimes (in ns) of HB and CB interactions (Section 3 of main text, corresponding to Figures 11-15). Total simulation time of 1000 ns. We indicate in "Bond" the atomic site of the designed new drug DBD-15-21-22 and the amino acid or ion involved in the HB or CB and the corresponding atoms in parentheses. Double labels indicate averaged values.

| Bond (drug/ion/amino acid (atom)) | KRAS-G12D |
|---|---|
| DBD15-21-22 (H12)-ASP12 (O1,O2) | 389 |
| DBD15-21-22 (H13)-GDP (O2B) | 970 |
| DBD15-21-22 (H1)-ASP57 (O2) | 932 |
| DBD15-21-22 (H11)-THR35 (O1) | 330 |
| DBD15-21-22 (H11)-ILE36 (O1) | 53 |
| DBD15-21-22 (H11)-GLU37 (O3) | 32 |
| DBD15-21-22 (N4)-Mg | 750 |
| DBD15-21-22 (N5)-Mg | 1000 |
| ASP57 (O1)-Mg | 1000 |
| SER17 (O2)-Mg | 964 |
| GDP (O1B)-Mg | 1000 |
| Water (O)-Mg | 931 |

#### 3 Supplementary Figures

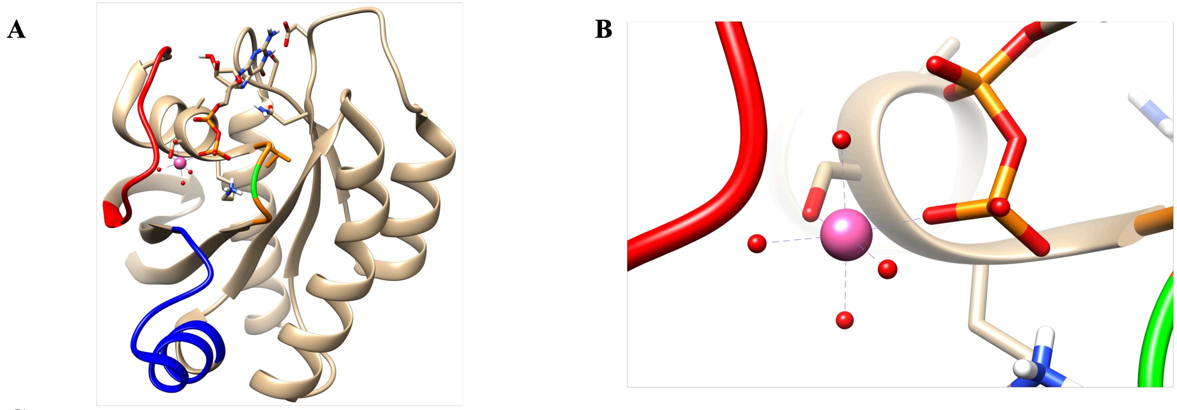


**C**

**Sequence of KRAS-WT**

10 20 30 40 50 60 70 80 90
MTEYKLVVVGAGGVGKSALTIQLIQNHFVDEYDPTIEDSYRKQVVIDGETCLLDILDTAGQEEYSAMRDQYMRTGEGFLCVFAINNTKSF
EDIHHYREQIKRVKDSEDVPMVLVGNKCDLPSRTVDTKQAQDLARSYGIPFIETSAKTRQGVDDAFYTLVREIRKHKEK
100 110 120 130 140 150 160

**Sequence of KRAS-G12C**

10 20 30 40 50 60 70 80 90
MTEYKLVVVGACGVGKSALTIQLIQNHFVDEYDPTIEDSYRKQVVIDGETCLLDILDTAGQEEYSAMRDQYMRTGEGFLCVFAINNTKSF
EDIHHYREQIKRVKDSEDVPMVLVGNKCDLPSRTVDTKQAQDLARSYGIPFIETSAKTRQGVDDAFYTLVREIRKHKEK
100 110 120 130 140 150 160

**Sequence of KRAS-G12D**

10 20 30 40 50 60 70 80 90
MTEYKLVVVGADGVGKSALTIQLIQNHFVDEYDPTIEDSYRKQVVIDGETCLLDILDTAGQEEYSAMRDQYMRTGEGFLCVFAINNTKSF
EDIHHYREQIKRVKDSEDVPMVLVGNKCDLPSRTVDTKQAQDLARSYGIPFIETSAKTRQGVDDAFYTLVREIRKHKEK
100 110 120 130 140 150 160

**Figure 1.** The initial structure and sequence of three KRAS isoforms: (A) SW-I (red), SW-II (blue), mutation site GLY12 (green); (B) Details of the octahedral structure of $Mg^{2+}$ in the crystal structure; (C)Sequence of KRAS-WT, KRAS-G12C and KRAS-G12D. G12 (green), C12 (yellow), D12 (cyan).

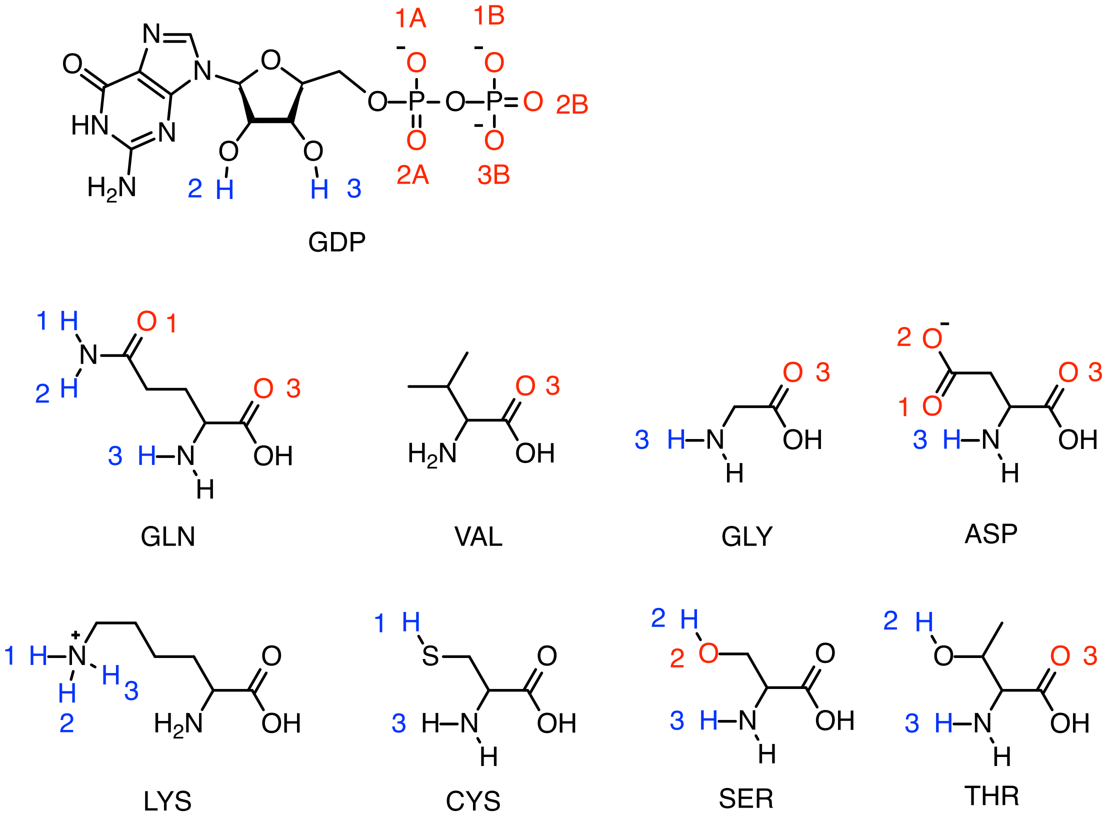


**Figure 2.** Sketches of GDP, and amino acid residues of KRAS protein mentioned in the text.

#### References

1. John, J. *et al.* Kinetic and structural analysis of the mg (2+)-binding site of the guanine nucleotide-binding protein p21h-ras. *Journal of Biological Chemistry* **268**, 923–929 (1993).

2. Bock, C. W., Kaufman, A. & Glusker, J. P. Coordination of water to magnesium cations. *Inorganic Chemistry* **33**, 419–427 (1994).

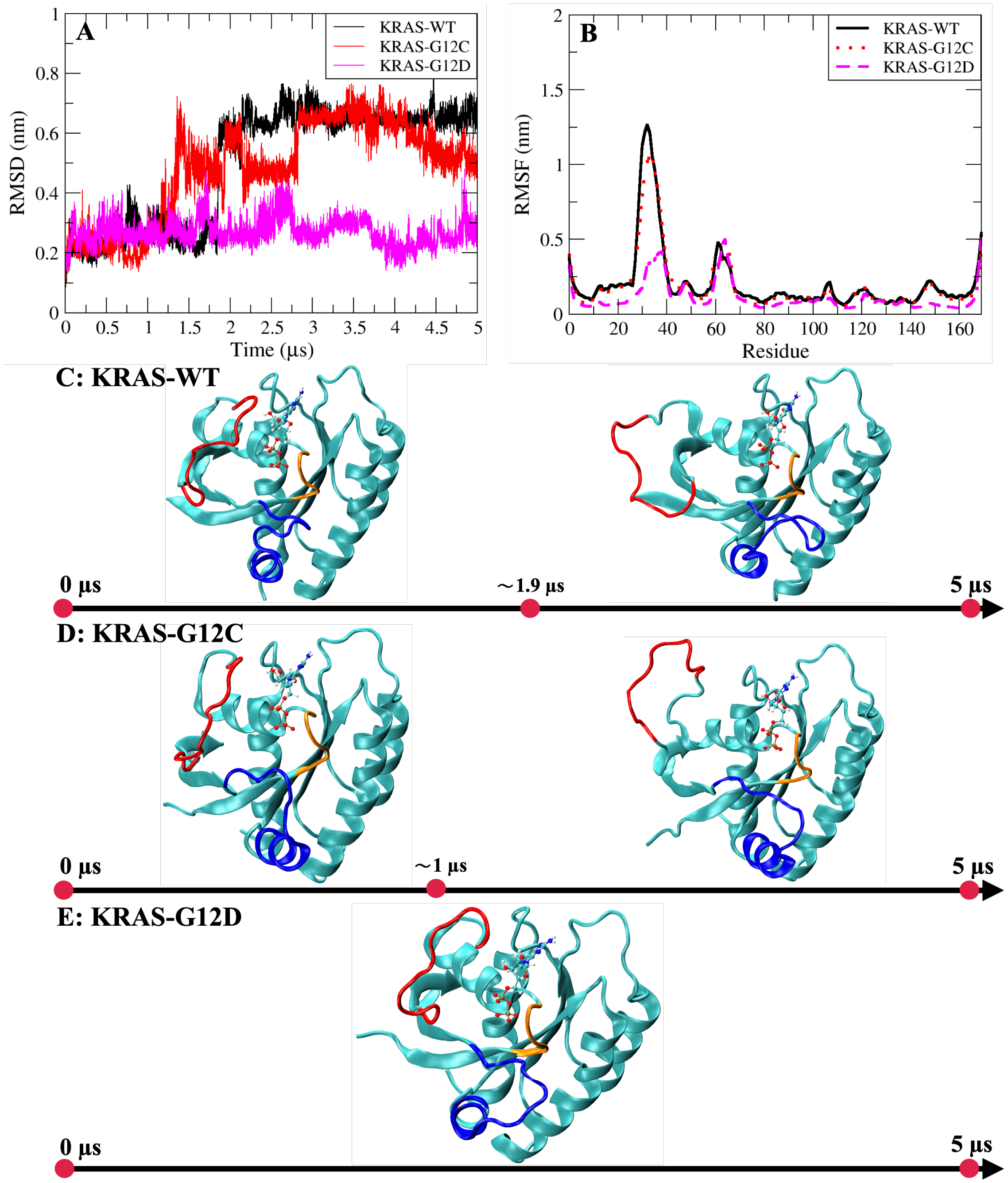


**Figure 3.** (A) Comparison of RMSD values of three KRAS proteins; (B) Comparison of RMSF values of three KRAS proteins; (C) The evolution of KRAS-WT conformational changes; (D) The evolution of KRAS-G12C conformational changes; (E) The evolution of KRAS-G12D conformational changes. Red: Switch-I; Blue: Switch-II; Orange: P-loop. Water and ions have been hidden.

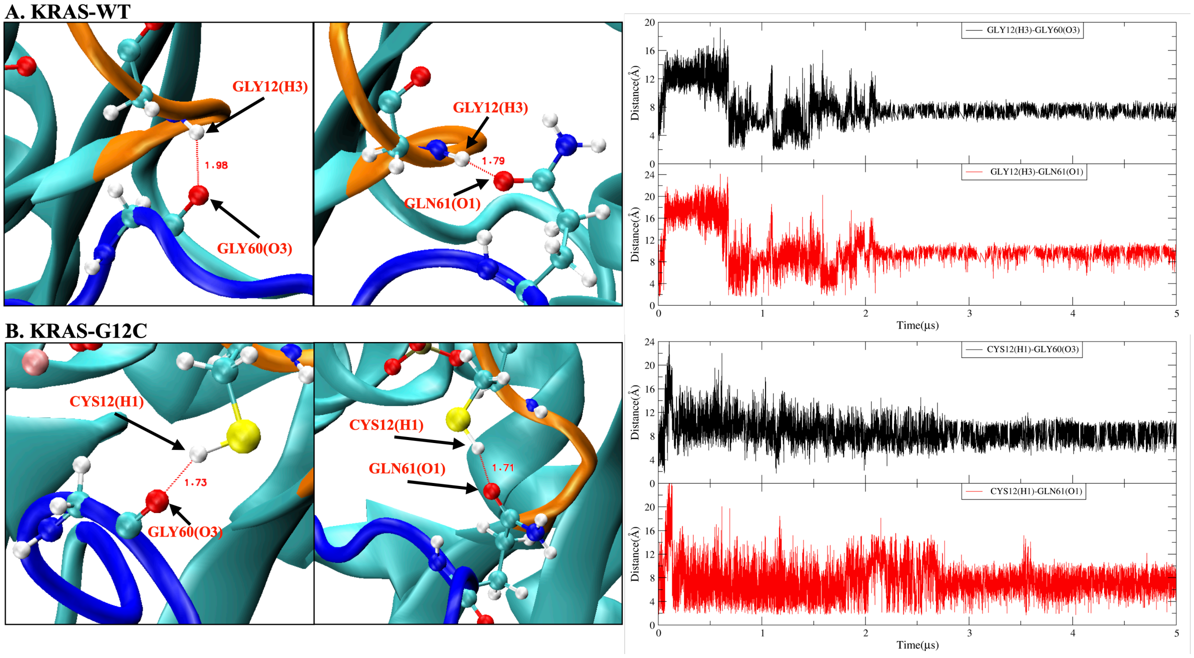


**Figure 4.** Analysis of hydrogen bond interaction between KRAS mutation site 12 and amino acid residues (GLY60, GLN61) of SW-II domain. (A) KRAS-WT case. Left side: representative snapshot of $HB_{GLY12(H3)-GLY60(O3)}$ and $HB_{GLY12(H3)-GLN61(O1)}$, right side: time evolution of atom-atom distance $d(t)$ corresponding to the left side; (B) KRAS-G12C case. Left side: representative snapshot of $HB_{CYS12(H1)-GLY60(O3)}$ and $HB_{CYS12(H1)-GLN61(O1)}$, right side: time evolution of atom-atom distance $d(t)$ corresponding to the left side.

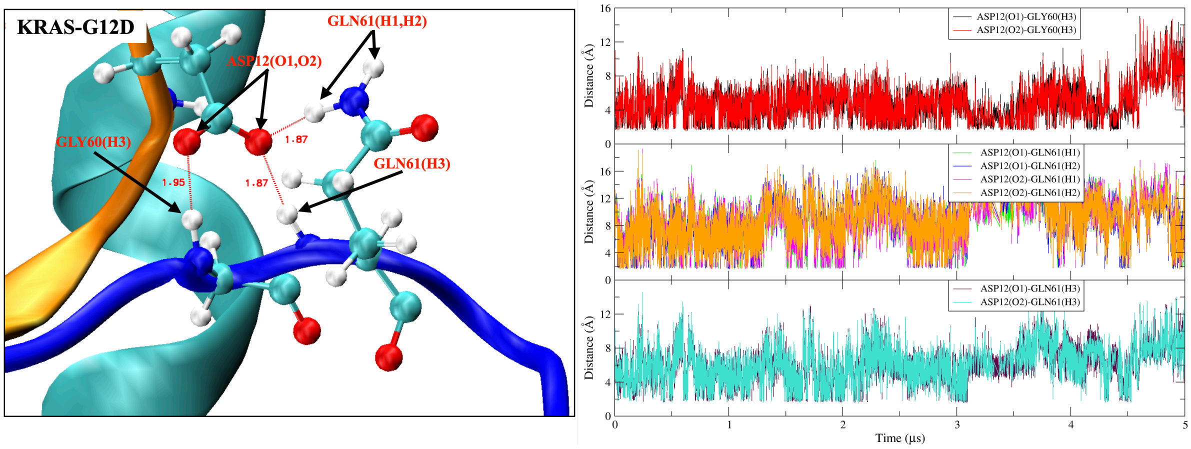


**Figure 5.** Analysis of hydrogen bond interaction between KRAS-G12D ASP12 residue and amino acid residues (GLY60, GLN61) of SW-II domain.(A) Representative snapshot of $HB_{ASP12(O1/O2)-GLY60(H3)}$, $HB_{ASP12(O1/O2)-GLN61(H1/H2)}$ and $HB_{ASP12(O1/O2)-GLN61(H3)}$; (B) time evolution of atom-atom distance $d(t)$ corresponding to (A).

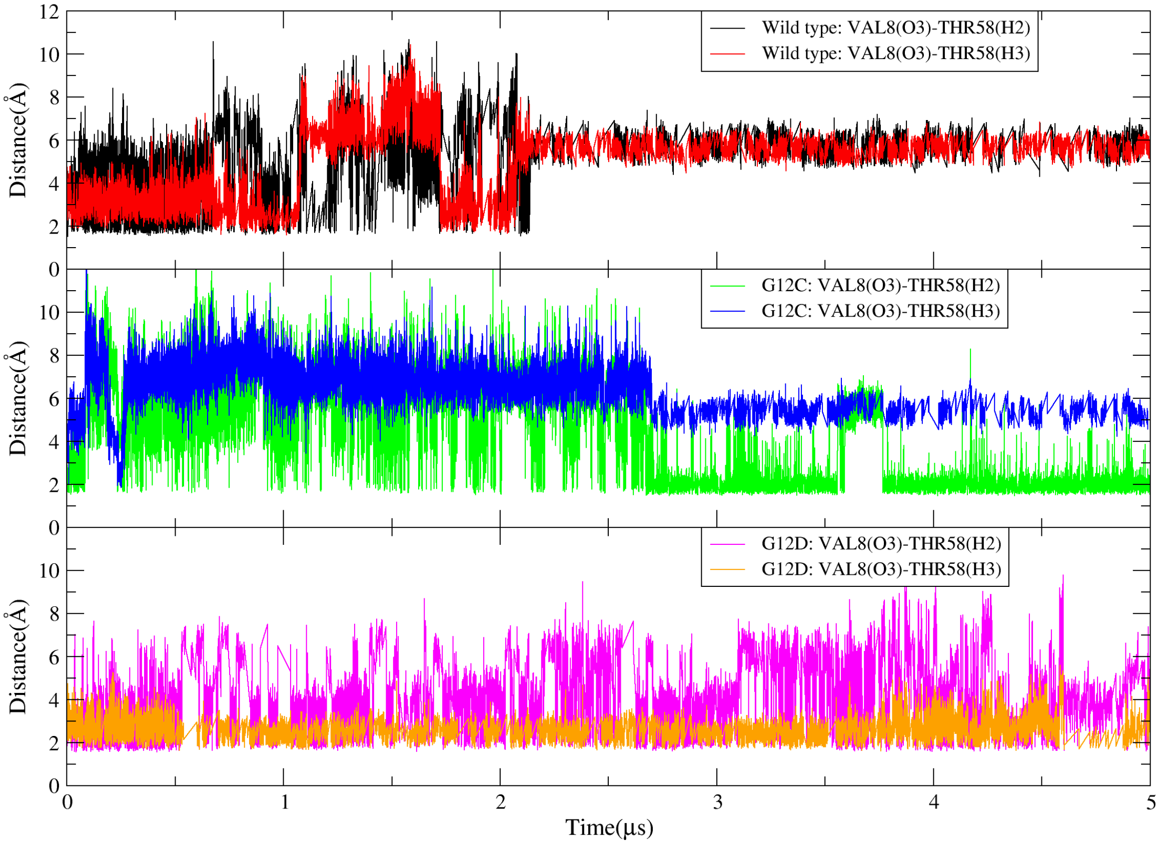


**Figure 6.** Atom-atom distance between oxygen atom (O3) of residue VAL8 and hydrogen atom (H2 and H3 of THR58) of KRAS SW-II.

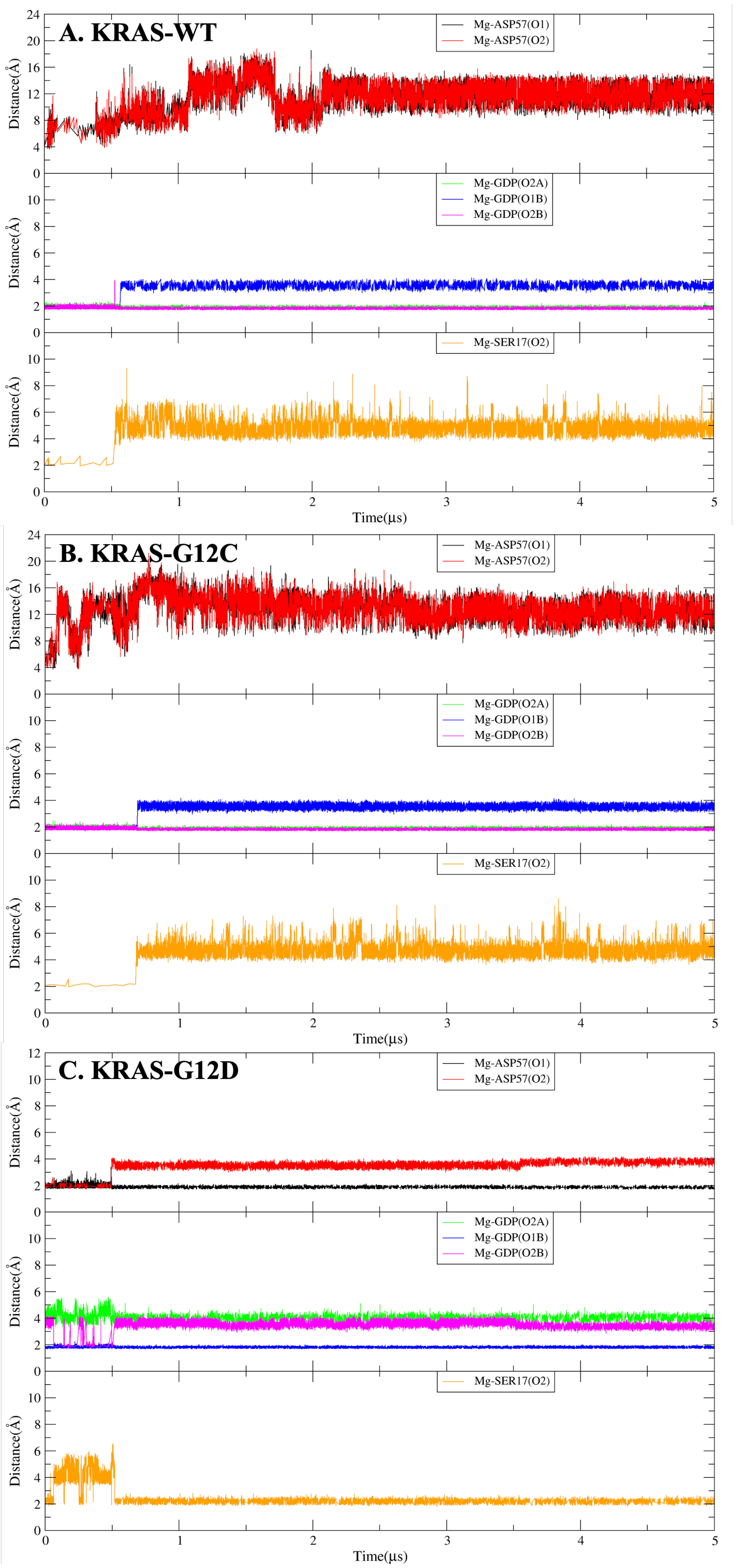


**Figure 7.** Coordination bonds analysis of $Mg^{2+}$ with related amino acid residues and GDP. (A) The $Mg^{2+}$ interaction with KRAS-WT-GDP.; (B) The $Mg^{2+}$ interaction with KRAS-G12C-GDP; (C) The $Mg^{2+}$ interaction with KRAS-G12D-GDP.

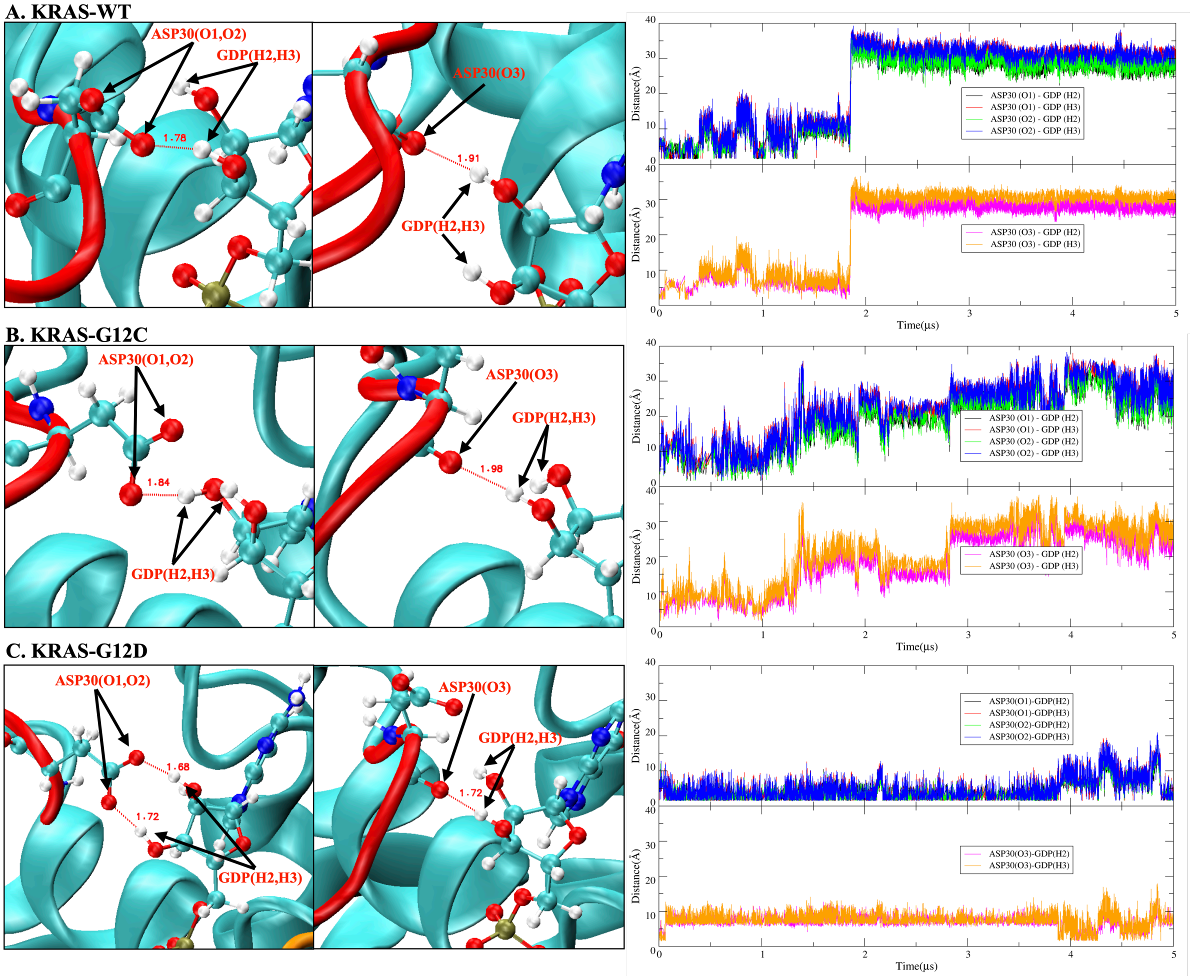


**Figure 8.** The hydrogen bond interaction between ASP30 of KRAS and GDP. (A) KRAS-WT case. Left side: representative snapshot of $HB_{ASP30(O1/O2)-GDP(H2/H3)}$ and $HB_{ASP30(O3)-GDP(H2/H3)}$, right side: time evolution of atom-atom distance $d(t)$ corresponding to the left side; (B) KRAS-G12C case. Left side: representative snapshot of $HB_{ASP30(O1/O2)-GDP(H2/H3)}$ and $HB_{ASP30(O3)-GDP(H2/H3)}$, right side: time evolution of atom-atom distance $d(t)$ corresponding to the left side; (C) KRAS-G12D case. Left side: representative snapshot of $HB_{ASP30(O1/O2)-GDP(H2/H3)}$ and $HB_{ASP30(O3)-GDP(H2/H3)}$, right side: time evolution of atom-atom distance $d(t)$ corresponding to the left side.

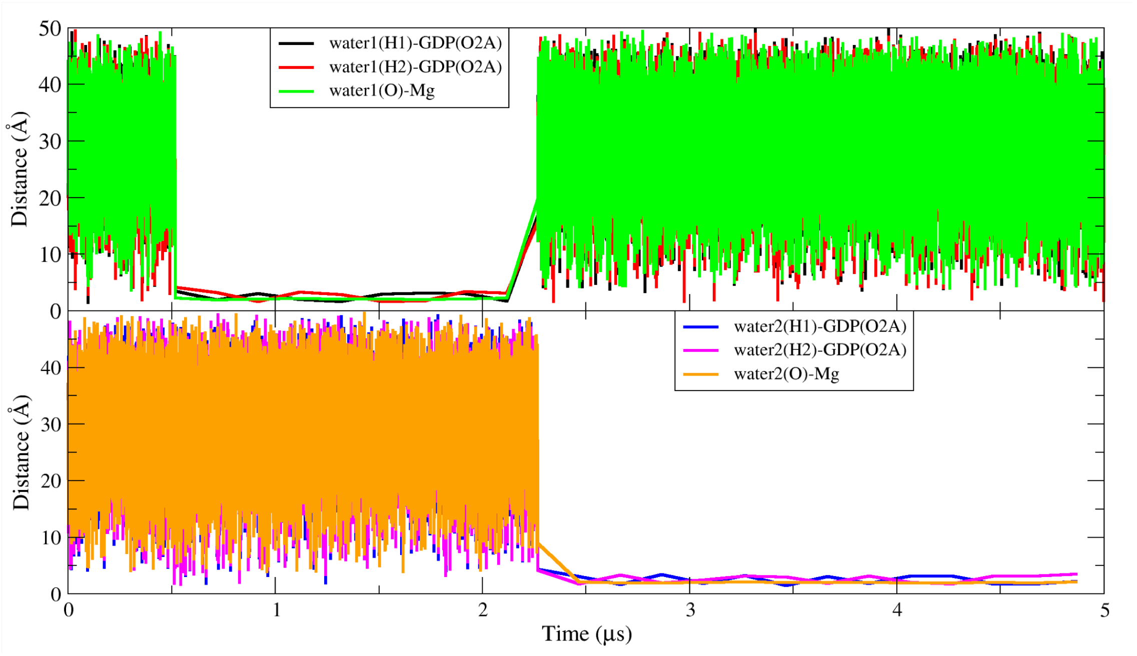


**Figure 9.** The dynamic change law of water molecules in the dynamic water pocket I.

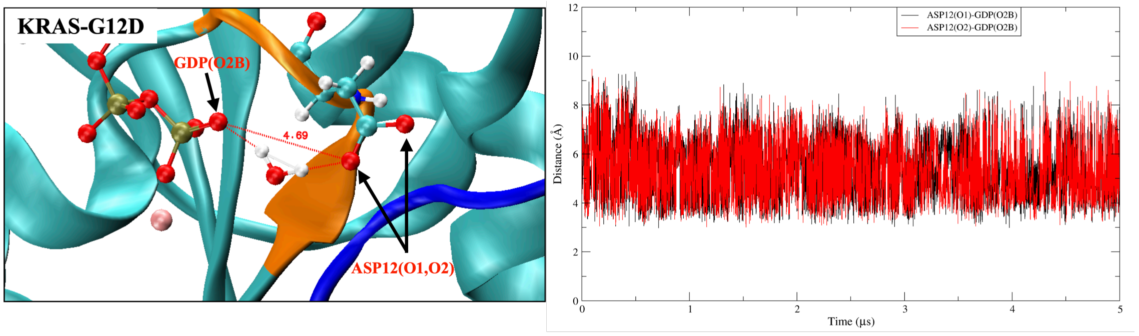


**Figure 10.** The dynamic water pocket II

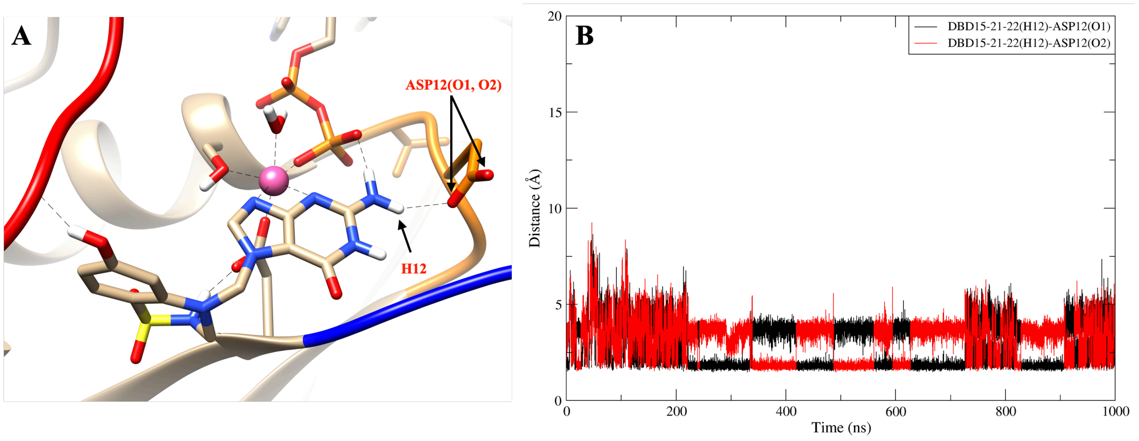


**Figure 11.** The hydrogen bond interaction between DBD15-21-22 (H12) and ASP12 (O1, O2). (A) Representative snapshot of $HB_{DBD15-21-22(H12)-ASP12(O1/O2)}$; (B) Distance between DBD15-21-22 (H12) and ASP12 (O1, O2).

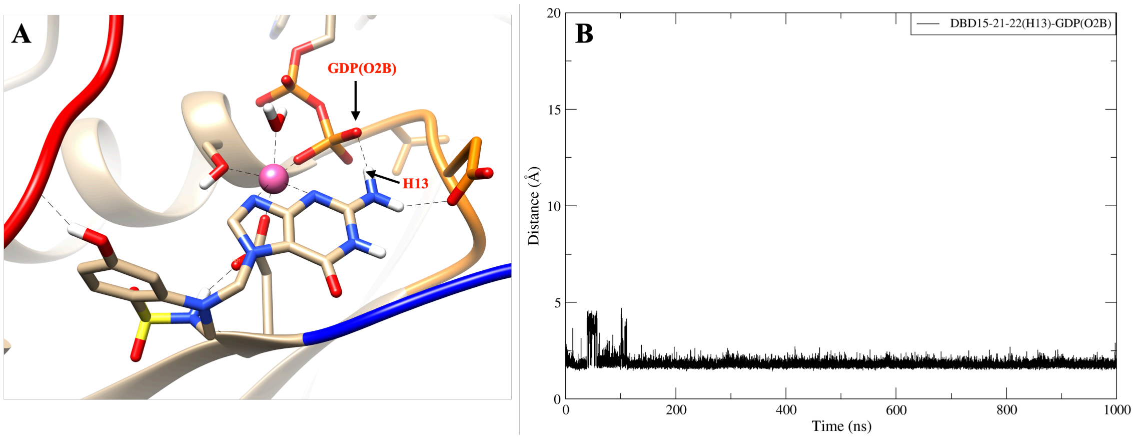


**Figure 12.** The hydrogen bond interaction between DBD15-21-22 (H13) and GDP (O2B). (A) Representative snapshot of $HB_{DBD15-21-22(H13)-GDP(O2B)}$; (B) Distance between DBD15-21-22 (H13) and GDP (O2B).

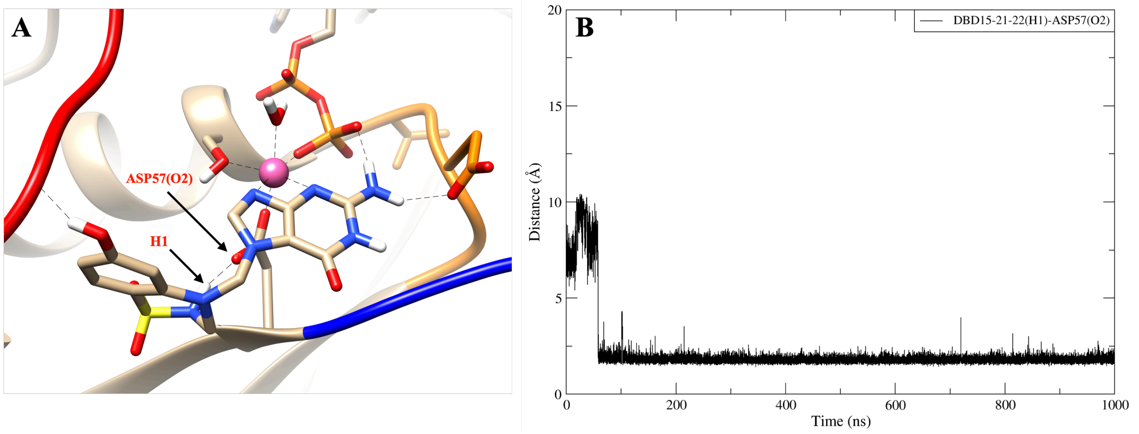


**Figure 13.** The hydrogen bond interaction between DBD15-21-22 (H1) and ASP57 (O2).(A) Representative snapshot of $HB_{DBD15-21-22(H1)-ASP57(O2)}$; (B) Distance between DBD15-21-22 (H1) and ASP57 (O2).

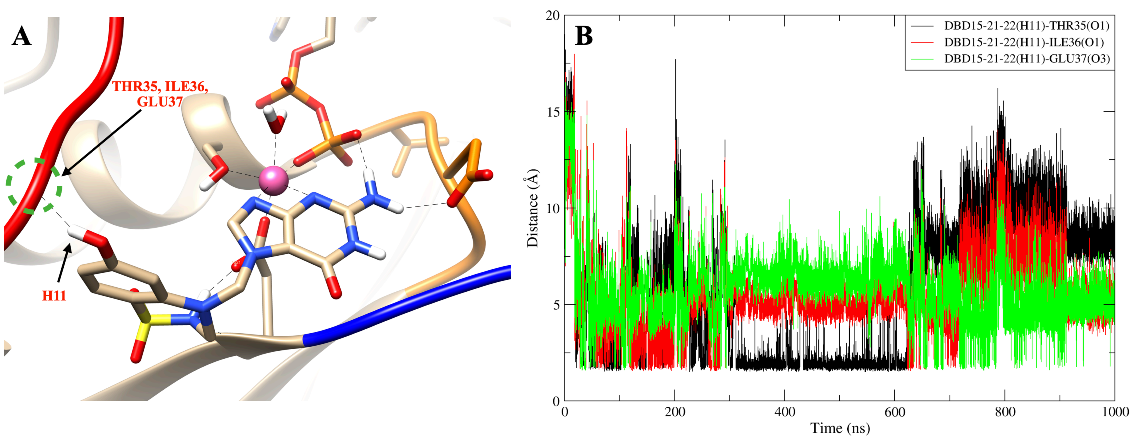


**Figure 14.** The DBD15-21-22 (H11) can form hydrogen bond interactions with THR35, ILE36 and GLU37. (A) Representative snapshot of DBD15-21-22 (H11) forming hydrogen bond interactions with THR35, ILE36 and GLU37; (B) Distance between DBD15-21-22 (H11) and three SW-I residues(THR35, ILE36, GLU37).

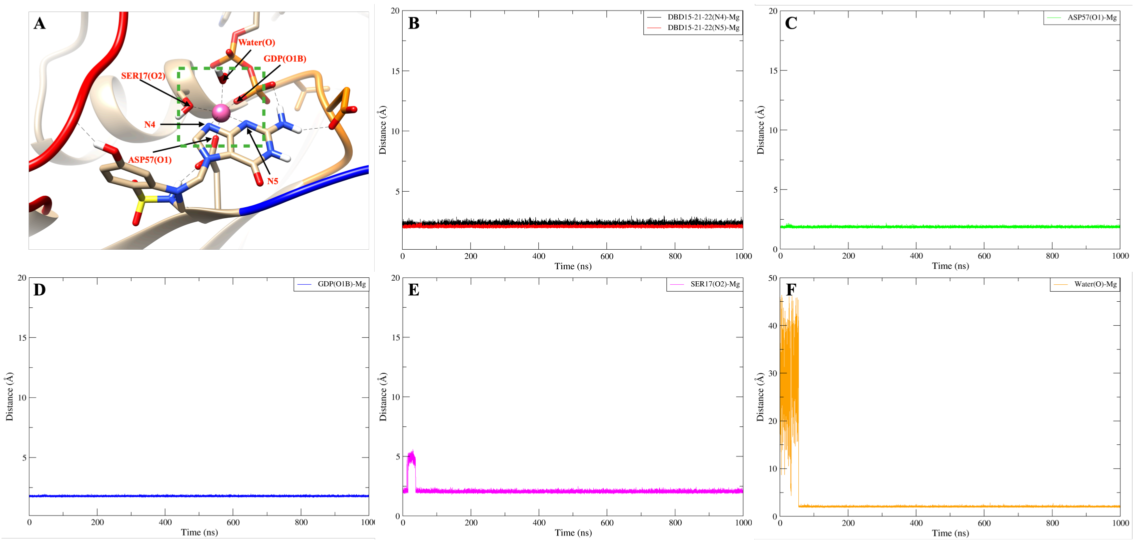


**Figure 15.** DBD15-21-22 binding on the dynamic water pocket I and stabilised the octahedral structure of $Mg^{2+}$. (A) Representative snapshot of DBD15-21-22 (N4,N5) form coordination bond with $Mg^{2+}$; (B) Distance between DBD15-21-22 (N4,N5) and $Mg^{2+}$; (C) Distance between ASP57 (O1) and $Mg^{2+}$; (D) Distance between GDP (O1B) and $Mg^{2+}$; (E) Distance between SER17(O2) and $Mg^{2+}$; (F) Distance between water molecular in position $H_2O-1$ and $Mg^{2+}$.

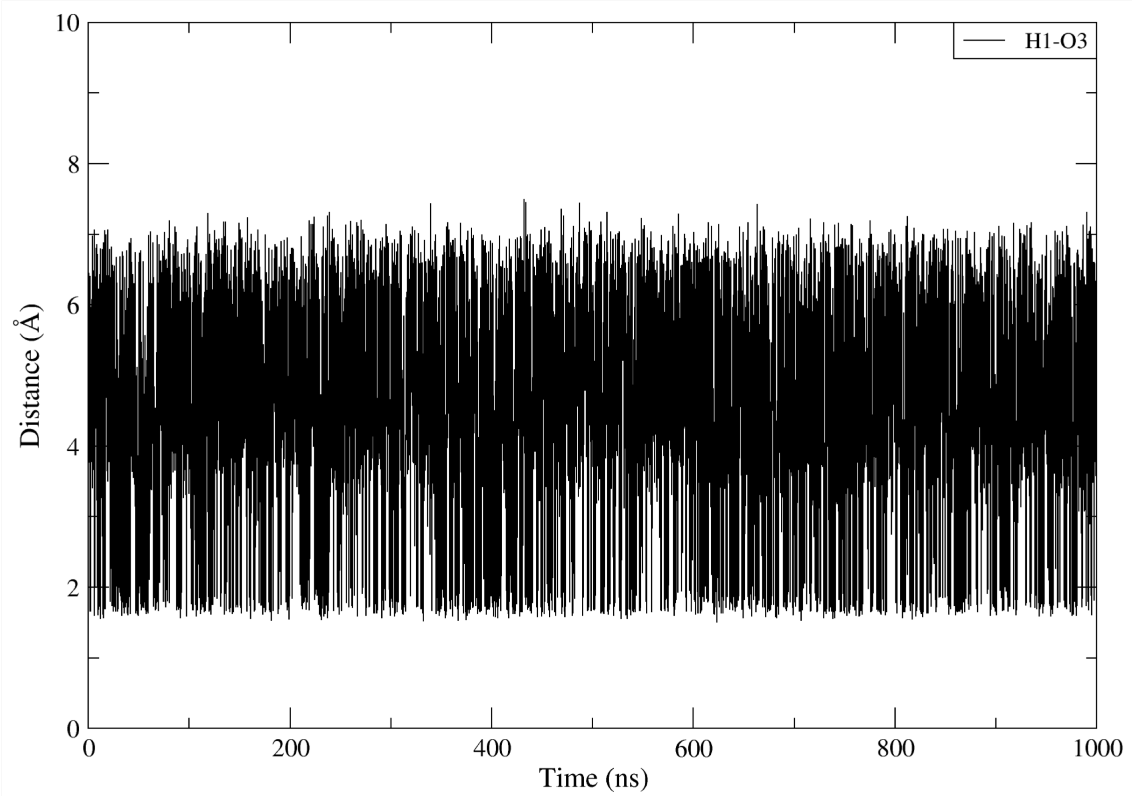


**Figure 16.** The distance between H1 and O3 atoms in DBD15-21-22.

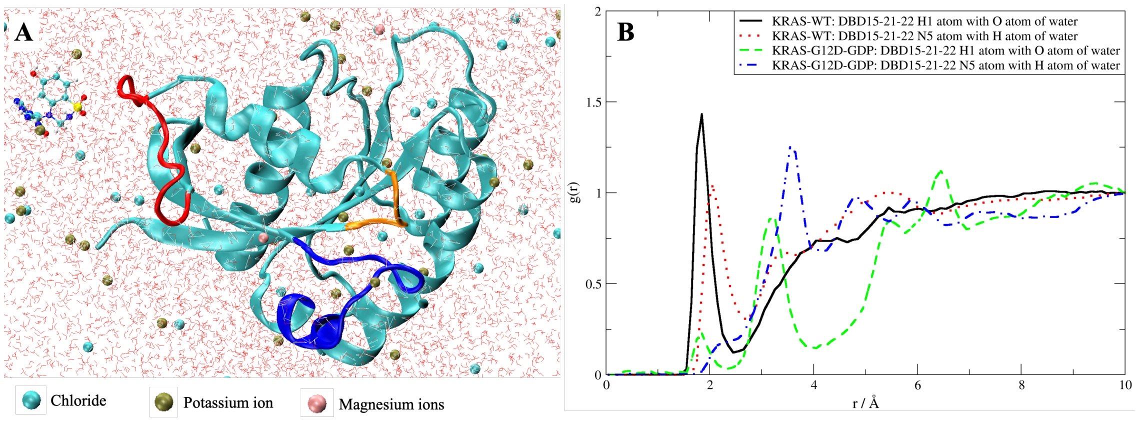


**Figure 17.** (A) Snapshot of DBD15-21-22 simulation with GDP/GTP free KRAS-WT and (B) RDF of DBD15-21-22 with water.

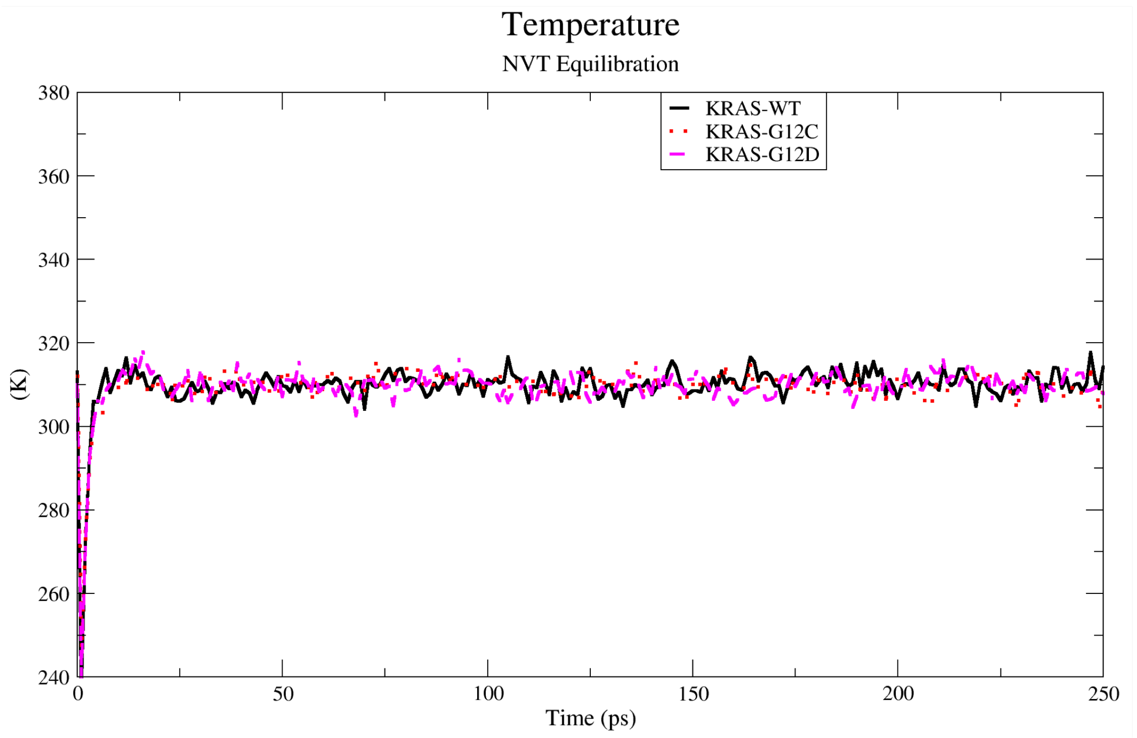


**Figure 18.** The three simulation systems (KRAS-WT, KRAS-G12C and KRAS-G12D) were well equilibrated before the production step.

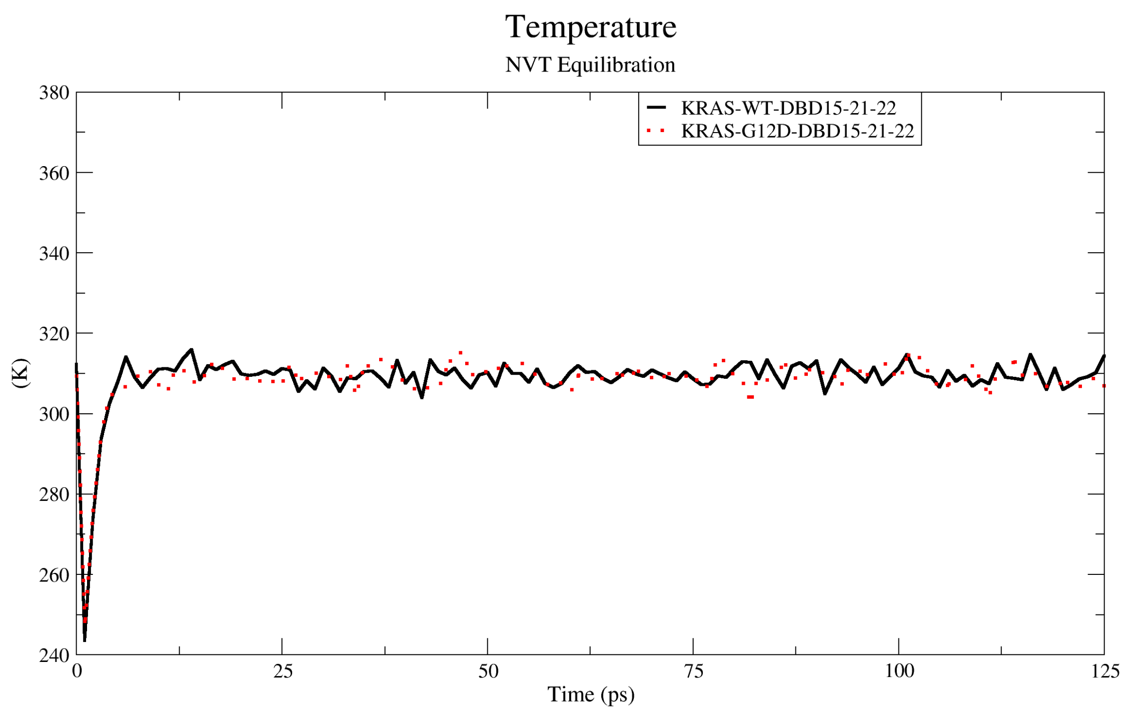


**Figure 19.** The two simulation systems (KRAS-WT with DBD15-21-22 and KRAS-G12D with DBD15-21-22) were well equilibrated before the production step.

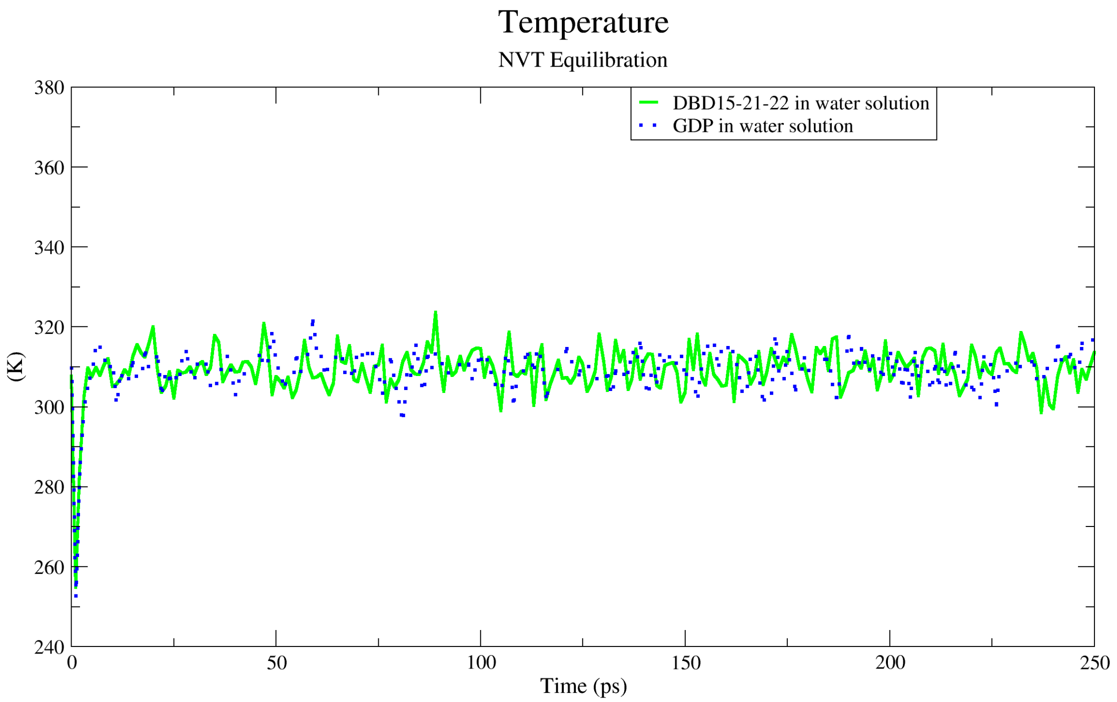


**Figure 20.** The two simulation systems (DBD15-21-22 in water solution, and GDP in water solution) were well equilibrated before the production step.
